# The 1p/19q co-deletion induces targetable and imageable vulnerabilities in glucose metabolism in oligodendrogliomas

**DOI:** 10.1101/2025.05.20.655097

**Authors:** Suresh Udutha, Georgios Batsios, Céline Taglang, Anne Marie Gillespie, Pavithra Viswanath

## Abstract

**Background:** The 1p/19q co-deletion is a hallmark of oligodendrogliomas. The goal of this study was to exploit metabolic vulnerabilities induced by the 1p/19q co-deletion for oligodendroglioma therapy and non-invasive imaging.

**Methods:** We used stable isotope tracing, mass spectrometry, and genetic and pharmacological approaches to interrogate [U-^13^C]-glucose metabolism in patient-derived oligodendroglioma models (SF10417, BT88, BT54, TS603, NCH612). We examined whether tracing [6,6’-^2^H]-glucose metabolism using deuterium metabolic imaging (DMI) provided an early readout of treatment response.

**Results:** The expression of the glycolytic enzyme enolase 1 (ENO1; chromosome 1p36.23) was reduced in patient-derived oligodendroglioma cells and patient biopsies due to the 1p/19q co-deletion and histone hypermethylation. Conversely, ENO2 was upregulated, an effect that was driven by mitogen-activated protein kinase (MAPK) signaling and ERK1-mediated phosphorylation and inactivation of the *CIC* transcriptional repressor in oligodendrogliomas. Genetic ablation of ENO2 or pharmacological inhibition using POMHEX inhibited proliferation with nanomolar potency but was not cytotoxic to oligodendroglioma cells or tumor xenografts. Mechanistically, ENO2 loss abrogated [U-^13^C]-glucose metabolism to lactate but shunted glucose towards biosynthesis of serine and purine nucleotides, an effect that was driven by phosphoglycerate dehydrogenase (PHGDH). Importantly, the PHGDH inhibitor D8 was synthetically lethal in combination with POMHEX, and the combination induced tumor regression *in vivo.* Furthermore, DMI of lactate production from [6,6’-^2^H]-glucose provided an early readout of response to combination therapy that preceded MRI-detectable alterations and reflected extended survival.

**Conclusions:** We have identified ENO2 and PHGDH as 1p/19q co-deletion-induced metabolic vulnerabilities in oligodendrogliomas and demonstrated that DMI reports on early response to therapy.

**KEY POINTS:**

- The 1p/19q co-deletion upregulates ENO2 in oligodendrogliomas.
- ENO2 inhibition inhibits glycolysis but upregulates serine and nucleotide biosynthesis via PHGDH.
- Combined inhibition of ENO2 and PHGDH is lethal, an effect that can be visualized by DMI.

**IMPORTANCE OF THE STUDY:** Oligodendrogliomas are devastating primary brain tumors with long-lasting and life-altering effects on physical and cognitive function. The presence of a 1p/19q co-deletion defines oligodendrogliomas. Here, using clinically relevant patient-derived models and patient tissue, we show that the 1p/19q co-deletion leads to loss of the glycolytic enzyme ENO1 and upregulation of ENO2 in oligodendrogliomas. This provides a unique therapeutic opportunity since most cells rely on ENO1 for glycolysis. Targeting ENO2 using the brain-penetrant inhibitor POMHEX abrogates glycolysis but redirects glucose toward serine and nucleotide biosynthesis, an effect that is driven by PHGDH, the rate-limiting enzyme for serine biosynthesis. Importantly, combined treatment with POMHEX and the PHGDH inhibitor D8 is synthetically lethal *in vitro* and *in vivo.* Furthermore, visualizing glucose metabolism using DMI provides an early readout of response to therapy that predicts extended survival *in vivo*. In summary, we have developed a unique integrated metabolic therapy and imaging approach for oligodendrogliomas.

## INTRODUCTION

Diffuse gliomas are the most common malignant primary brain tumors in adults^1^. These tumors are now classified by their molecular features into 3 entities: glioblastoma (GBM, grade 4), oligodendroglioma (ODG, grades 2 and 3), and astrocytoma (AC, grades 2, 3, or 4)^2^. Although ODGs and ACs are indolent in growth relative to GBMs and confer a better prognosis, they tend to afflict younger patients (median age of ∼40 vs. ∼70 years) and cause debilitating seizures^1,3^. As such, ODG and AC patients are likely to be at the peak of their careers, socially and economically active, engaged in raising families, and their diagnosis has negative socioeconomic ramifications^1,3^.

ODGs and ACs harbor a mutation in isocitrate dehydrogenase 1 or, rarely, 2 (IDHm), while GBMs are IDH wild-type^4^. IDHm converts the housekeeping metabolite α-ketoglutarate to D-2-hydroxyglutarate (D-2HG), which inhibits α-ketoglutarate-dependent histone and DNA demethylases to induce epigenetic alterations that drive tumorigenesis^4^. Although IDHm is the earliest disease-defining oncogenic event in these tumors, secondary lineage-defining alterations and tertiary oncogenic alterations are needed for tumor formation^2,5^. For instance, ACs are driven by IDHm in concert with secondary lineage-defining alterations in *TP53* or *ATRX* and tertiary alterations in *PDGFRA* and *MET*^2,5^. In contrast, ODGs are driven by IDHm combined with a 1p/19q co-deletion and recurrent somatic mutations in the residual alleles of *CIC* (located on chromosome 19q) and *FUBP1* (located on chromosome 1p)^2,5–7^. Although the precise role of the 1p/19q co-deletion in orchestrating ODG growth is unclear, the observation that loss-of-function mutations in the residual alleles of CIC and FUBP are common in ODGs points to a tumor suppressive role for CIC and FUBP1^6^. In addition to these secondary alterations, ODGs often harbor tertiary alterations in the MAPK, PI3K/Akt, and KRAS pathways^7^. Of note, patient-derived ODG cells that harbor only IDHm, 1p/19q co-deletion, and CIC without additional tertiary alterations do not form xenografts in animals, suggesting that the activation of these pathways is essential for tumor formation *in vivo*^5^.

Standard treatment for ODG and AC patients includes maximal safe surgical resection, radiation, and alkylating chemotherapy (temozolomide or the combination of procarbazine, lomustine/CCNU, and vincristine)^8^. However, patients inevitably experience tumor recurrence and succumb to the disease, a phenomenon that is attributed to the fact that diffuse gliomas are highly infiltrative and extend far beyond the boundaries of the radiographically visible lesion. Furthermore, both radiation and chemotherapy are associated with side effects such as myelosuppression, neuropathy, infertility, and neurocognitive deficits that negatively impact survival and quality of life in this relatively young population^8^. Recently, the IDHm inhibitor vorasidenib, which suppresses D-2HG production and extends progression-free survival, was approved for the treatment of ODG and AC patients. While undeniably significant, not all patients respond to vorasidenib, and even in patients with responsive tumors, the gain in progression-free survival is only ∼18 months^9^.

Since the 1p/19q co-deletion is a molecular hallmark of ODGs^2,6^, unraveling the biological consequences of the 1p/19q co-deletion can lead to minimally toxic, precision therapies for ODG patients. One potential strategy is to target the isoform of a housekeeping gene that is lost due to the 1p/19q codeletion^10,11^. For instance, enolase is essential for the conversion of 2-phosphoglycerate to phosphoenolpyruvate during glycolysis. Enolase is encoded by 3 isoforms, i.e., the ubiquitously expressed ENO1 (located on 1p36.23), brain-specific ENO2 (12p13.31), and muscle-specific ENO3 (17p13.2). ENO1 loss confers sensitivity to ENO2 inhibition in a small subset of glioblastomas (∼1-5%) that lose ENO1 due to 1p36 deletions^11^. Importantly, POMHEX is a potent ENO2-specific inhibitor that halts the growth of ENO1-deleted glioblastomas *in vivo*^10,11^.

MRI methods such as contrast-enhanced MRI and T2/FLAIR are the mainstay for monitoring disease progression and treatment response in ODG patients^12^. However, ODGs often do not enhance on contrast-enhanced MRI and are notoriously difficult to visualize by MRI^13^. As a result, current response assessment guidelines require physicians to compare MRI scans separated by ≥8 weeks to determine whether a patient is responding to therapy^13,14^. This inability to quickly and reliably monitor response to therapy results in ineffective treatment for patients with progressive disease and, conversely, unnecessary toxicity for patients with responsive disease. Importantly, it also hampers the proper assessment of the efficacy of novel agents in clinical trials^15^.

Magnetic Resonance Spectroscopy (MRS) is an MR-compatible method of interrogating the magnetic resonance of tissue metabolites with a concentration >1 mM *in vivo*^16^. ^1^H-MRS, which interrogates the magnetic resonance of protons, provides a readout of metabolite abundance and has been used to detect D-2HG in glioma patients^16,17^. However, unambiguous D-2HG detection by ^1^H-MRS is complicated by its complex spectral pattern and overlap with other metabolites such as glutamate, glutamine, and γ-aminobutyric acid^18^. ^1^H-MRS also needs complex acquisition sequences, is susceptible to magnetic field inhomogeneity, and the presence of lipid and water suppression artifacts^18^. ^2^H-MRS or deuterium metabolic imaging (DMI) is a novel method of monitoring the metabolism of ^2^H-labeled substrates and, thereby, gaining a readout of metabolic activity (as opposed to abundance) *in vivo*^19–21^. We recently showed that [6,6′-^2^H]-glucose metabolism to [3,3′-^2^H]-lactate provides a readout of tumor burden in preclinical AC models *in vivo*^20^. Importantly, the feasibility of clinical translation of DMI has been established in normal volunteers and GBM patients^19,22–24^.

The goal of this study was to determine whether ENO1 loss due to the 1p/19q co-deletion confers a therapeutic opportunity in ODGs. Our studies indicate that ENO2 is upregulated in ODG cells and patient tissue in a MAPK pathway-driven manner. Genetic ablation of ENO2 or treatment with POMHEX abrogates glycolysis but redirects glucose toward the biosynthesis of serine and pyrimidine nucleotides, an effect that is mediated via upregulation of the rate-limiting enzyme for serine synthesis i.e., PHGDH. Importantly, inhibiting both ENO2 and PHGDH is synthetically lethal and induces tumor shrinkage *in vivo*, an effect that can be visualized within a few days of treatment by DMI.

## MATERIALS AND METHODS

Detailed methods are provided in the Supplementary Information.

### Cell culture

BT54, BT88, and BT142 cells were obtained from Dr. Luchman, the SF10417 and SF10602 lines from Dr. Costello, and TS603 cells from Dr. Venneti. The NCH612 cell line was obtained from Dr. Christel Herold-Mende from Heidelberg University and engineered to express luciferase for bioluminescence (BLI) imaging. NPCs were purchased from ATCC. Cells were cultured as described earlier^20,21,25^ (see Supplementary Methods).

### Patient biopsies

Patient biopsies were obtained from the UCSF Brain Tumor Center Biorepository in compliance with the written informed consent policy. Biopsy use was approved by the Committee on Human Research at UCSF and research was approved by the Institutional Review Board at UCSF according to ethical guidelines established by the U.S. Common Rule.

### Gene silencing and overexpression

ENO2 and PHGDH were silenced in ODG cells using AUM*silence^TM^* antisense oligonucleotides (ASOs; 5 μM). In each case, 2 different ASOs were used along with a non-targeting control. The ASOs were added to the cell culture medium at a concentration of 5 μM. Silencing was confirmed by QPCR at 72 h. Two non-overlapping Accell siRNA sequences (Dharmacon) were used to silence MEK1 or ERK1, or AMPK in ODG cells. Gene expression was confirmed by QPCR. The QuikChange II mutagenesis kit (Agilent) was used to generate the S173A CIC mutant in a pCMV6 expression vector (Origene, #RC215209).

### IC50

The Lumit® hKi-67 (Promega) was used to quantify live cell proliferation. Briefly, 10,000 cells/well were seeded in a 96-well plate and treated with a serial dilution of POMHEX or D8. Alternately, cells were treated with 5 μM non-targeting (NT) ASO, or ASO targeted against ENO2 or PHGDH. After 72 h, Ki67 expression was measured according to the manufacturer’s instructions. IC50 was calculated by non-linear regression. To assess synergy, 10,000 cells/well were seeded in a 96-well plate and treated with a 5-point serial dilution of POMHEX or D8. Live cell proliferation was measured using the Lumit® hKi-67 assay. Bliss synergy scores were calculated using SynergyFinder (version 3.0)^26^. For all functional assays, cells were treated with POMHEX or D8 at IC50 for 72 h. For the combination, cells were treated with a combination of drugs at doses that correspond to ≥90% inhibition based on the dose-response matrix from the synergy studies. Trametinib was used at a concentration of 50 nM.

### Flow cytometry

Cell cycle progression was measured by staining live cells for DNA content using the Vybrant™ DyeCycle™ Green stain (Invitrogen). Cells were analyzed by flow cytometry on a MACSQuant 10 analyzer. To measure doubling time, cells were seeded in a 96-well plate at 10,000 cells per well and treated as described above. After 4 days, the cell number was measured using the MACSQuant 10 analyzer. The doubling time was calculated using the following formula: doubling time = [4 days × (ln2)] / [ln (day 4 cell count/day 0 cell count)]. To quantify apoptosis in tumor tissue, single cell suspensions were depleted of mouse cells, stained with a 1:50 dilution of the anti-annexin V-PE antibody, washed, and annexin V+ cells were measured using a MACSQuant 10 flow cytometer. The PLA was performed as described in the supplementary methods^27^.

### Stable isotope tracing

Cells were treated as needed and incubated in media containing 25 mM [U-^13^C]-glucose for 72 h. Metabolites were extracted using methanol/water and analyzed by liquid chromatography-mass spectrometry (LC-MS)^28^. For *in vivo* studies, mice bearing intracranial BT88 or SF10417 tumors were treated as described below. At day 7, mice were intravenously injected with a bolus of 502.2 mg/kg of [U-^13^C]-glucose followed by continuous infusion at 50.2 mg/kg/min for 30 min^28,29^. Tumor and contralateral normal brain tissue were snap frozen, and metabolites were extracted using methanol/water^28^.

### LC-MS

LC-MS was performed using a Vanquish Ultra High-performance LC coupled to an Orbitrap ID-X mass spectrometer^28^. Metabolites were separated using a Luna 3 NH2 column (150 mm x 2.1 mm, 3 μm, Phenomenex). High-resolution MS data was acquired using a full scan method alternating between positive and negative polarities. Peak areas were quantified using TraceFinder, and fractional ^13^C enrichment was calculated after correcting for natural abundance using Escher Trace^28^.

### In vivo studies

All studies were approved by the University of California, San Francisco Institutional Animal Care and Use Committee (IACUC). ODG cells (5-6x10^5^ cells in 3 μL) were intracranially injected into the cortex of SCID mice (female, 5-6 weeks)^20,21,28^. Once tumors were visible on MRI or BLI, this timepoint was considered D0, and mice were randomized. Mice with BT88 tumors were treated with vehicle (saline), POMHEX (50 mg/kg), or the combination of POMHEX and D8 (50 mg/kg each) every day for 5 days per week via intraperitoneal injection. For the NCH612 model, mice were treated with vehicle (saline), POMHEX (10 mg/kg), or the combination of POMHEX (10 mg/kg) and D8 (15 mg/kg) daily for 5 days per week via oral gavage. In both cases, mice were treated until they needed to be euthanized per IACUC guidelines, or the tumor was no longer visible on MRI. Tumor volume was determined by T2-weighted MRI using a Bruker 3T or 9.4T scanner^20,21,28^. Radiance was quantified on a Spectral Instruments Imaging (ATX) after intraperitoneal injection of 150 mg/kg D-luciferin.

### DMI

Studies were performed on a Bruker 3T or 9.4T scanner using a 16 mm ^2^H surface coil in combination with a quadrature ^1^H coil (40 mm inner diameter). For spatial localization of lactate production, a 2D chemical shift imaging (CSI) sequence was used (3T: TE/TR = 1.04/265.89 ms, FOV = 30 x 30 x 8 mm^3^, complex points=128 points, spectral width=2.5 kHz, NA = 30, temporal resolution=8 minutes 30 s, nominal voxel size = 112.5 μL; 9.4T: TE/TR = 1.104/591.824 ms, FOV = 30x30x5 mm^3^, 512 points, spectral width = 2.5 kHz, NA = 8, temporal resolution=8 minutes 30 s, nominal voxel size = 70.3 μL. Data was analyzed using in-house MATLAB codes^20,21^.

### Statistical analysis

All experiments were performed on a minimum of 3 biological replicates (n≥3), and the results were expressed as mean ± standard deviation. Statistical significance was assessed in GraphPad Prism 10 using a two-way ANOVA or two-tailed Welch’s t-test, with p<0.05 considered significant. Analyses were corrected for multiple comparisons using Tukey’s or Dunnett’s method, where applicable. Survival was quantified using Kaplan-Meier analysis. * indicates p<0.05, ** indicates p<0.01, *** indicates p<0.001 and **** indicates p<0.0001.

### Data availability

Data is available from the corresponding author upon reasonable request.

## RESULTS

### ENO2 is uniquely upregulated in patient-derived ODG models and patient biopsies

First, we quantified ENO1 expression in patient-derived ODG and AC models. Since proliferating ODG cells closely resemble neural progenitor cells (NPCs) in their expression signatures^30^, we examined NPCs as added controls. As shown in Fig. 1A-1C, ENO1 mRNA and protein were significantly downregulated in ODG (SF10417, BT88, BT54, TS603, NCH612) cells relative to AC (BT142, SF10602) and NPCs. In contrast, ENO2 mRNA and protein were significantly upregulated in ODG cells relative to AC and NPC (Fig. 1D-1F). To validate the clinical relevance of our results, we examined biopsies from ODG or AC patients. As shown in Fig. 1G-1J, ENO1 mRNA and protein were significantly reduced, while ENO2 was significantly elevated in ODG biopsies relative to AC. These results are consistent with TCGA data from the GLASS consortium showing that ENO1 mRNA is significantly reduced in IDHm 1p/19 co-deleted tumors (ODGs) relative to IDHm non-co-deleted tumors (ACs; Fig. 1K). Importantly, interrogation of pan-cancer TCGA data indicated that ENO2 expression was highest in ODGs relative to all other tumor types (Supplementary Fig. S1).

**Figure 1.**
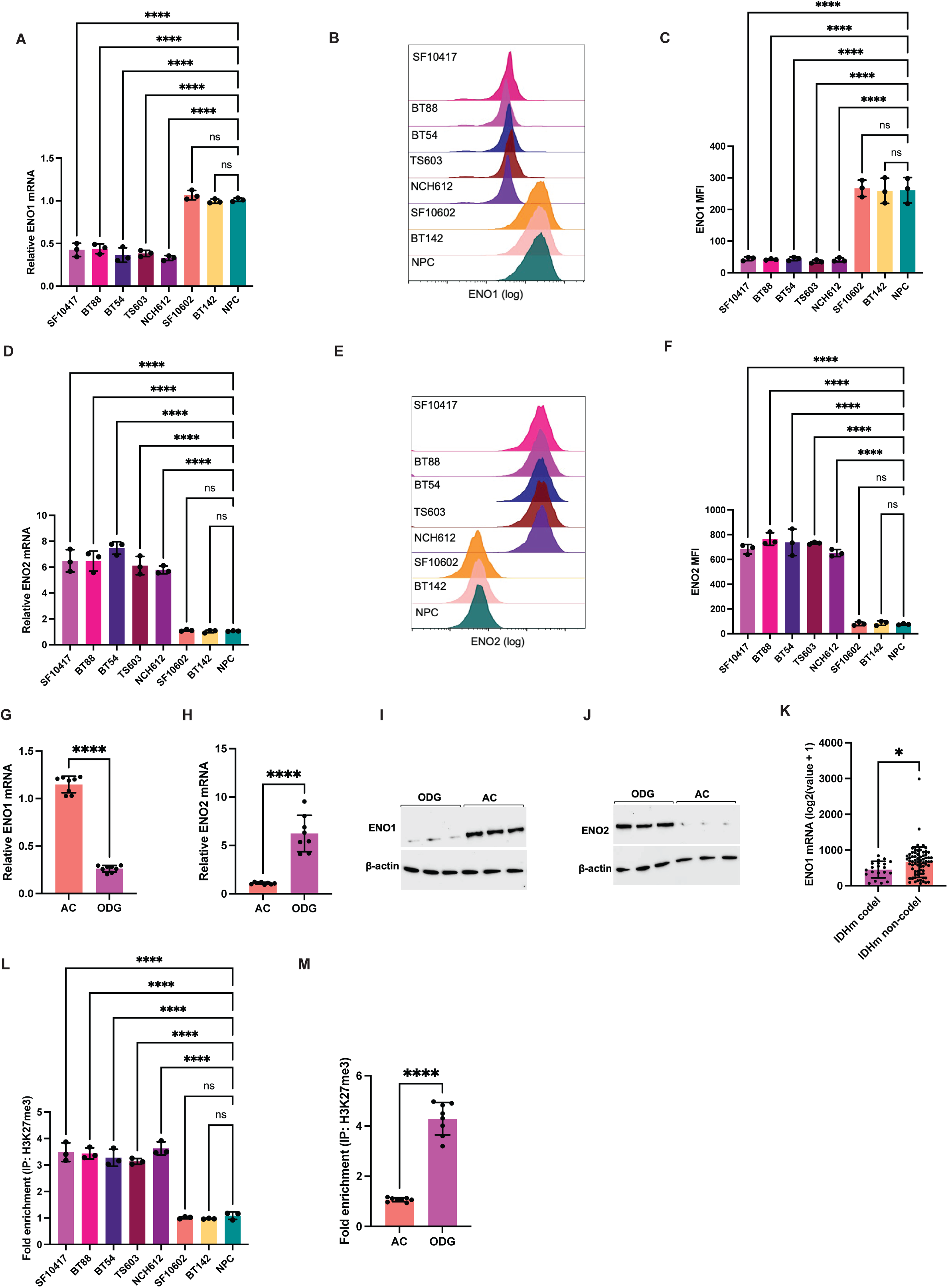
ENO1 is downregulated while ENO2 is upregulated in ODGs. **(A)** ENO1 mRNA abundance in patient-derived ODG (SF10417, BT88, BT54, TS603, NCH612), AC (SF10602, BT142), and NPCs. Representative flow cytometric histograms of ENO1 staining **(B)** and quantification of ENO1 staining intensity **(C)** in patient-derived ODG, AC, and NPCs. **(D)** ENO2 mRNA abundance in patient-derived ODG, AC, and NPCs. Representative flow cytometric histograms of ENO2 staining **(E)** and quantification of ENO2 staining intensity **(F)** in patient-derived ODG, AC, and NPCs. ENO1 **(G)** and ENO2 **(H)** mRNA in biopsies from ODG or AC patients. Western blots for ENO1 **(I)** and ENO2 **(J)** expression in ODG and AC biopsies. **(K)** The abundance of ENO1 mRNA in tissue from ODG (IDH mutant co-deleted (ODG) and non-co-deleted (AC) tumors in the GLASS consortium database. **(L)** H3K27me3 enrichment at the ENO1 promoter in patient-derived ODG, AC, and NPCs. **(M)** H3K27me3 enrichment at the ENO1 promoter in biopsies from ODG or AC patients.

The 1p/19 co-deletion is expected to lead to the loss of one allele of ENO1 and, thereby, reduce ENO1 expression. However, our data above indicated that ENO1 expression was almost completely lost in ODG cells and patient biopsies. IDHm is known to suppress the expression of glycolytic genes, including lactate dehydrogenase A and the monocarboxylate transporter 4, via histone hypermethylation^31,32^. As shown in Fig. 1L-1M, the enrichment of histone H3K27me3 at the ENO1 promoter was significantly elevated in both patient-derived ODG cells and patient biopsies relative to AC. These studies indicate that ENO1 expression is significantly reduced due to the 1p/19q co-deletion and histone hypermethylation, while, conversely, ENO2 expression is significantly upregulated in ODGs relative to ACs and normal cells.

### The MAPK pathway upregulates ENO2 in ODGs

Next, we examined the molecular mechanism by which ENO2 is upregulated in ODGs. Previous studies indicate that ODGs harbor tertiary alterations that activate the MAPK and PI3K/Akt pathways^5,7^. Since these signaling pathways are known to reprogram glucose metabolism in tumor cells^33^, we questioned whether they regulate ENO2 expression in ODGs. We first confirmed that levels of phosphorylated ERK1 and AKT, which are biomarkers of the MAPK and PI3K/Akt pathways, respectively, were higher in patient-derived ODG models (SF10417, BT88, BT54, TS603, NCH612) relative to AC (SF10602, BT142) and NPCs (Fig. 2A-2B and Supplementary Fig. S2A-S2B). Phosphorylated ERK1 and Akt were also higher in ODG patient tissue relative to AC (Fig. 2C). We then treated patient-derived ODG cells with a panel of 15 signaling pathway inhibitors and quantified ENO2 expression by ELISA. As shown in Fig. 2D, inhibition of the MAPK pathway downregulated ENO2 expression with maximal effect following MEK1/2 (mirdametinib, trametinib) or ERK1/2 (temuterkib, MK-83853) inhibition. To further confirm these results, we examined the effect of silencing MEK1 or ERK1 in ODG cells (Supplementary Fig. S2C-S2D). As shown in Fig. 2E, silencing MEK1 or ERK1 abrogated ENO2 expression in ODG cells. These results indicate that the MAPK pathway upregulates ENO2 expression in ODGs.

**Figure 2.**
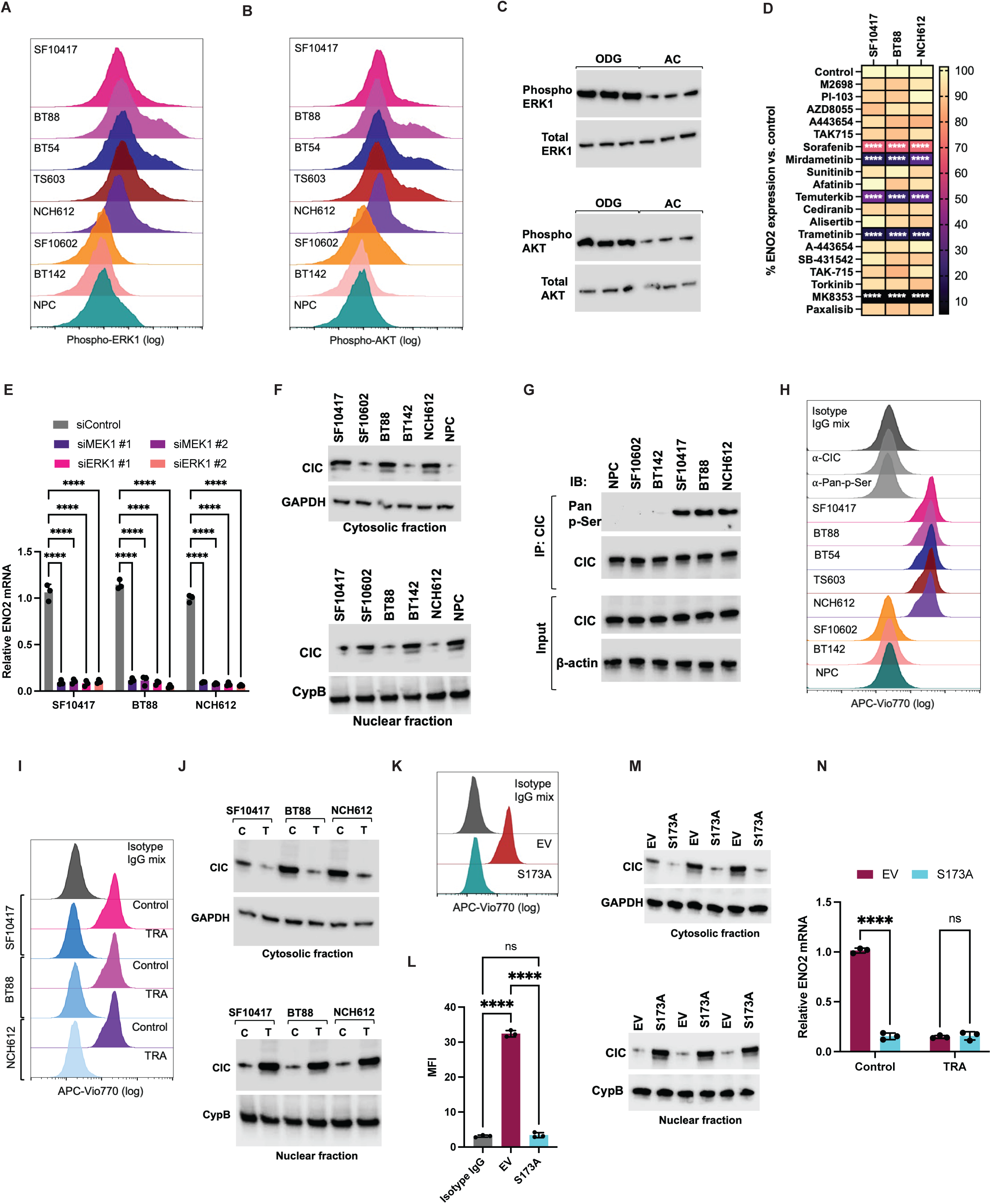
ERK1-driven phosphorylation of CIC derepresses ENO2 expression in ODGs. Representative flow cytometric histograms for phospho-ERK1 **(A)** and phospho-AKT **(B)** in patient-derived ODG (SF10417, BT88, BT54, TS603, NCH612), AC (SF10602, BT142), and NPCs. **(C)** Western blots for phospho-ERK1 (top panel) and phospho-AKT (bottom panel) in biopsies from ODG or AC patients. **(D)** Heatmap of the effect of signaling pathway inhibitors on ENO2 expression in ODG (SF10417, BT88, NCH612) cells. **(E)** Effect of silencing MEK1 or ERK1 using 2 non-overlapping siRNA sequences on ENO2 mRNA abundance in ODG cells. **(F)** Western blots for CIC expression in the cytosol (top panel) vs. nucleus (bottom panel) in patient-derived ODG (SF10417, BT88, NCH612), AC (SF10602, BT142), and NPCs. GAPDH was used as the loading control for the cytosolic fraction, and cyclophilin B (CypB) for the nuclear fraction. **(G)** Western blots demonstrating CIC phosphorylation in NPC, AC (SF10602, BT142) and ODG (SF10417, BT88, NCH612) cells. Lysates were immunoprecipitated with an anti-CIC (α-CIC) antibody and probed for phosphorylation using an anti-Pan-phosphoserine (α-Pan-p-Ser) antibody. IP refers to immunoprecipitation, and IB refers to immunoblotting. The top panel shows the IP, and the bottom panel shows the input for IB at 10%. **(H)** Representative flow cytometric histograms of the PLA for CIC phosphorylation in patient-derived ODG (SF10417, BT88, BT54, TS603, NCH612), AC (SF10602, BT142), and NPCs. Cells were treated with a mixture of the mouse α-CIC and rabbit α-Pan-p-Ser antibodies. Cells treated with the mixture of the corresponding isotype IgG antibodies (referred to as Isotype IgG mix) or with either primary antibody alone were used as negative controls. **(I)** Representative flow cytometric histograms of the PLA for CIC phosphorylation in ODG cells treated with vehicle (control) or trametinib (TRA). SF10417 cells treated with the mixture of the corresponding isotype IgG antibodies (Isotype IgG mix) were used as controls. **(J)** Western blots for CIC expression in the cytosol (top panel) vs. nucleus (bottom panel) in patient-derived ODG (SF10417, BT88, NCH612) cells treated with vehicle (C) or trametinib (T). GAPDH was used as the loading control for the cytosolic fraction, and cyclophilin B (CypB) for the nuclear fraction. **(K)** Representative flow cytometric histograms of the PLA for CIC phosphorylation in SF10417 cells expressing the empty vector (EV) or S173A mutant CIC. **(L)** Quantification of CIC phosphorylation by the PLA in SF10417 cells expressing the empty vector (EV) or S173A mutant CIC. **(M)** Western blots for CIC expression in the cytosol (top panel) vs. nucleus (bottom panel) in SF10417 cells expressing the empty vector (EV) or S173A mutant CIC. **(N)** ENO2 mRNA in SF10417 cells expressing the empty vector (EV) or S173A mutant CIC in the presence or absence of trametinib.

### ERK1-driven phosphorylation inactivates CIC and derepresses ENO2 expression in ODGs

We then questioned whether the MAPK pathway upregulates ENO2 in a manner that is linked to the 1p/19q co-deletion. Although the precise mechanism by which the 1p/19q co-deletion contributes to oligodendrogliomagenesis is unknown, the observation that the residual allele for CIC, which is located on chromosome 19q and is partially lost due to the 1p/19q codeletion, is mutated in >70% of ODGs, points to a role for CIC as a tumor suppressor^6^. Functionally, CIC is a high-mobility group (HMG)-box transcriptional repressor that downregulates the expression of MAPK pathway targets, in part via the recruitment of histone deacetylases to the promoters of target genes^34,35^. Previous studies indicate that activated MAPK signaling results in ERK1-mediated phosphorylation of CIC at serine 173 (S173), which can lead to ubiquitin-mediated degradation or cytosolic CIC sequestration^35^. In both cases, the expression of CIC target genes is activated.

SF10417 and BT88 cells harbor a *CIC* mutation, while NCH612 is CIC wild-type^36–38^. We confirmed that CIC mRNA was detectable, albeit significantly reduced, in all patient-derived ODG cells relative to ACs and NPCs (Supplementary Fig. S2E). Strikingly, the CIC protein was predominantly sequestered in the cytosol in ODG cells (SF10417, BT88, NCH612), unlike ACs (SF10602, BT142) and NPCs, where it was largely localized to the nucleus (Fig. 2F). Immunoprecipitation followed by immunoblotting confirmed that CIC was phosphorylated in ODG cells but not NPCs (Fig. 2G). We also confirmed these results using a proximity ligation assay (PLA), which allows *in situ* detection of endogenous protein modifications^27^ (Fig. 2H and Supplementary Fig. S2F). Importantly, inhibiting MAPK signaling by treatment with trametinib abrogated CIC phosphorylation and caused cytosolic sequestration of CIC in ODG cells (Fig. 2I-2J and Supplementary Fig. S2G). These results suggest that ERK1 phosphorylates CIC and prevents its nuclear localization in ODGs.

To determine whether CIC is phosphorylated at S173 and assess its role in ENO2 expression, we expressed FLAG-tagged mutant CIC S173A, which is non-phosphorylatable, in SF10417 cells (Supplementary Fig. S2H). The S173A mutation blocked CIC phosphorylation and increased nuclear localization in SF10417 cells (Fig. 2K-2M). Importantly, the S173A mutation downregulated ENO2 expression and rendered it insensitive to MAPK inhibition (Fig. 2N). These results identify CIC as a transcriptional repressor of ENO2 expression and indicate that ERK1-driven CIC phosphorylation at S173 derepresses ENO2 expression in ODGs.

### Genetic or pharmacological inhibition of ENO2 inhibits ODG proliferation and cell cycle progression

Given the unique upregulation of ENO2, we questioned whether targeting ENO2 had therapeutic potential for ODGs. To this end, we silenced ENO2 using ASOs in patient-derived ODG cells (Supplementary Fig. S3A). As controls, we examined AC cells and NPCs. As shown in Fig. 3A-3C, ENO2 silencing inhibited proliferation as measured by Ki67 expression and blocked cell cycle progression at the S phase specifically in ODG but not AC cells or NPCs. To further confirm these results, we examined the effect of pharmacologically inhibiting ENO2 using the potent, ENO2-specific inhibitor POMHEX^10,11^. As shown in Fig. 3D, POMHEX inhibited the viability of ODG cells with IC50 values of 35-50 nM. In contrast, the IC50 for ACs and NPCs was >5-20 μM, pointing to the exquisite dependence of ODG cells on ENO2 activity (Fig. 3D). Similar to ENO2 silencing, treatment with POMHEX significantly inhibited ODG proliferation and cell cycle progression (Fig. 3E-3F). There was no effect on ACs or NPCs (Fig. 3F). Of note, neither ENO2 silencing nor POMHEX induced cell death in ODG cells (Supplementary Fig. S3B-S3C). These results indicate that targeting ENO2 has a cytostatic effect on ODG proliferation.

**Figure 3.**
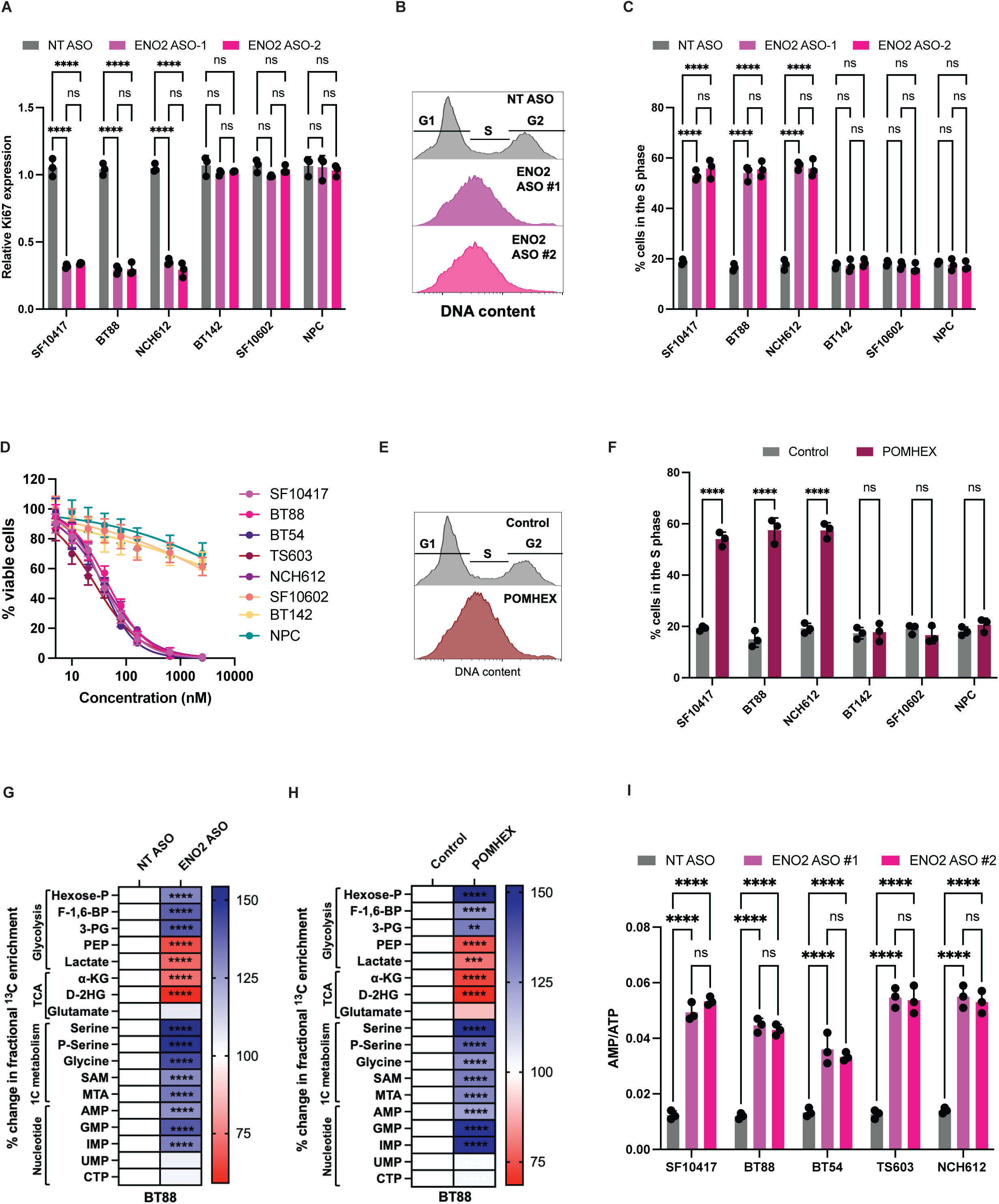
Targeting ENO2 inhibits ODG proliferation in a cytostatic manner. **(A)** Effect of silencing ENO2 using 2 non-overlapping ASO sequences on the proliferation of patient-derived ODG (SF10417, BT88, NCH612), AC (SF10602, BT142), or NPCs. Proliferation was measured by quantifying Ki67 expression. Cells treated with a non-targeting (NT) ASO were used as controls. Representative flow cytometric histograms from SF10417 cells **(B)** and quantification **(C)** of the effect of ENO2 silencing on cell cycle progression in patient-derived ODG, AC, or NPCs. **(D)** Dose-response curves showing the effect of POMHEX on the proliferation of patient-derived ODG, AC, or NPCs. Representative flow cytometric histograms from SF10417 cells **(E)** and quantification **(F)** of the effect of POMHEX on cell cycle progression in patient-derived ODG, AC, or NPCs. Effect of ENO2 silencing **(G)** or POMHEX **(H)** on [U-^13^C]-glucose metabolism in BT88 cells. **(I)** Effect of ENO2 silencing on the AMP/ATP ratio in patient-derived ODG cells. For panels G and H, Hexose-P refers to glucose-6-phosphate or fructose-6-phosphate; F-1,6-BP refers to fructose-1,6-bisphosphate; 3-PG is 3-phosphoglycerate; PEP is phosphoenolpyruvate; α-KG is α-ketoglutarate; P-serine is phosphoserine; SAM is S-adenosylmethionine; MTA is methylthioadenosine.

### Targeting ENO2 rewires [U-^13^C]-glucose metabolism via PHGDH in ODGs

Next, we used [U-^13^C]-glucose and liquid chromatography-mass spectrometry (LC-MS) to trace the effect of ENO2 silencing on glucose metabolism in ODG cells. ENO2 silencing significantly reduced fractional ^13^C labeling of glycolytic (phosphoenolpyruvate, lactate) and tricarboxylic acid (TCA) cycle (α-ketoglutarate, D-2HG) metabolites downstream of ENO2 (Fig. 3G). In contrast, ENO2 silencing increased fractional ^13^C labeling of metabolites associated with serine synthesis and one-carbon metabolism (3-phosphoglycerate, serine, phosphoserine, glycine, S-adenosylmethionine, methylthioadenosine) and purine nucleotide biosynthesis (AMP, GMP) in ODG cells (Fig. 3G). ENO2 silencing did not alter ^13^C labeling of pyrimidine nucleotides (UMP, CTP) from [U-^13^C]-glucose in ODG cells (Fig. 3G). We observed similar effects following pharmacological inhibition of ENO2 using POMHEX in ODG cells (Fig. 3H). Consistent with the inhibition of glycolysis, ATP abundance was significantly reduced following ENO2 silencing or inhibition (Supplementary Fig. S3D-S3E), leading to an elevated AMP/ATP ratio (Fig. 3I and Supplementary Fig. S3F).

We then investigated the mechanism by which ENO2 loss shunts glucose towards serine biosynthesis in ODG cells. PHGDH is the rate-limiting enzyme in serine synthesis from glucose, and its expression and/or activity is often upregulated in cancer^39^. As shown in Fig. 4A-4D, both ENO2 silencing and POMHEX upregulated PHGDH mRNA and protein in patient-derived ODG cells. Since AMPK is a central regulator of cell metabolism^40^, and our studies indicated that ENO2 loss increased the AMP/ATP ratio (see Fig. 3I and Supplementary Fig. 3F), which is known to activate AMPK via phosphorylation^40^, we questioned whether AMPK played a role in regulating PHGDH in ODGs. Indeed, as shown in Fig. 4E-4F, both ENO2 silencing and POMHEX resulted in AMPK phosphorylation in ODG cells. Importantly, silencing AMPK normalized PHGDH expression and activity in ENO2-ASO-treated cells to levels observed in controls (Fig. 4G-4J). Collectively, our studies indicate that targeting ENO2 in ODGs downregulates glycolysis but shunts glucose towards serine biosynthesis in an AMPK- and PHGDH-dependent manner.

**Figure 4.**
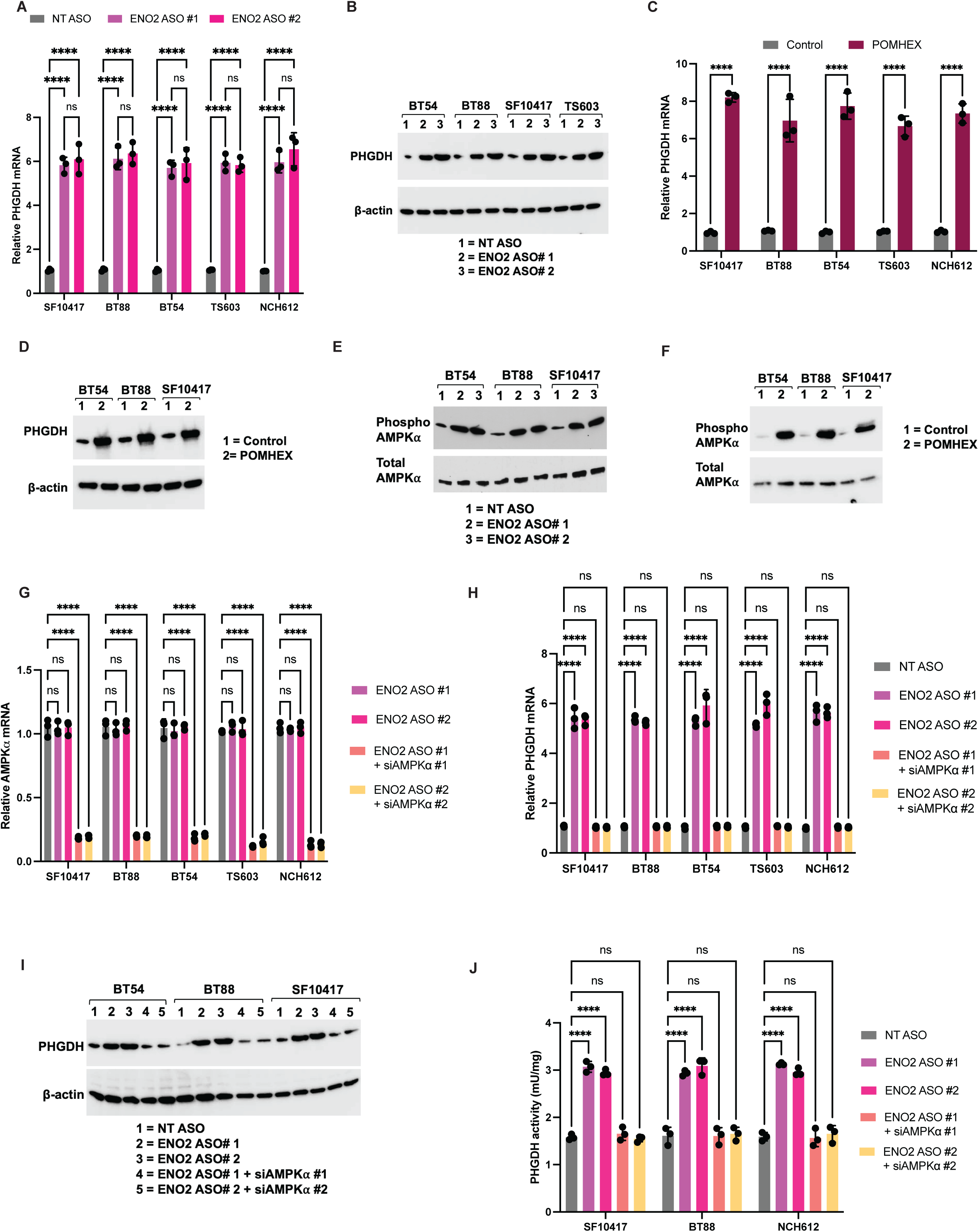
ENO2 silencing upregulates PHGDH in an AMPK-dependent manner in ODGs. Effect of ENO2 silencing on PHGDH mRNA **(A)** and protein **(B)** in patient-derived ODG cells. β-actin was used as the loading control for panel B. Effect of POMHEX on PHGDH mRNA **(C)** and protein **(D)** in patient-derived ODG cells. β-actin was used as the loading control for panel D. Western blots for activated AMPKα (phosphorylated at T172) in patient-derived ODG cells treated with ENO2 ASOs **(E)** or POMHEX **(F)**. Cells treated with the NT ASO or vehicle (DMSO) were used as controls. Total AMPKα was used as the loading control. **(G)** Verification of AMPKα silencing by QPCR for AMPKα mRNA in patient-derived ODG cells treated with the NT ASO, ENO2 ASO alone, or ENO2 ASOs in combination with AMPKα ASOs. Effect of silencing AMPKα on PHGDH mRNA **(H)**, protein **(I)**, and activity **(J)** in ODG cells treated with the NT ASO, ENO2 ASO alone, or ENO2 ASOs in combination with AMPKα ASOs.

### Targeting PHGDH is synthetically lethal in combination with ENO2 in ODG cells

Next, we examined the therapeutic potential of targeting both ENO2 and PHGDH in ODGs. Silencing both ENO2 and PHGDH using ASOs (Supplementary Fig. S4A-S4B) abrogated [U-^13^C]-glucose metabolism via glycolysis (lactate), the TCA cycle (α-ketoglutarate, D-2HG), serine and one-carbon metabolism (serine, phosphoserine, glycine, S-adenosylmethionine, methylthioadenosine), and purine nucleotide biosynthesis (AMP, GMP) in BT88 cells (Fig. 5A). Importantly, combined ENO2 and PHGDH silencing was synthetically lethal and resulted in massive apoptosis relative to non-targeted controls, in contrast to silencing ENO2 alone or PHGDH alone (Fig. 5B). This effect was specific to ODGs since combined ENO2 and PHGDH silencing did not induce apoptosis in ACs or NPCs (Fig. 5B).

**Figure 5.**
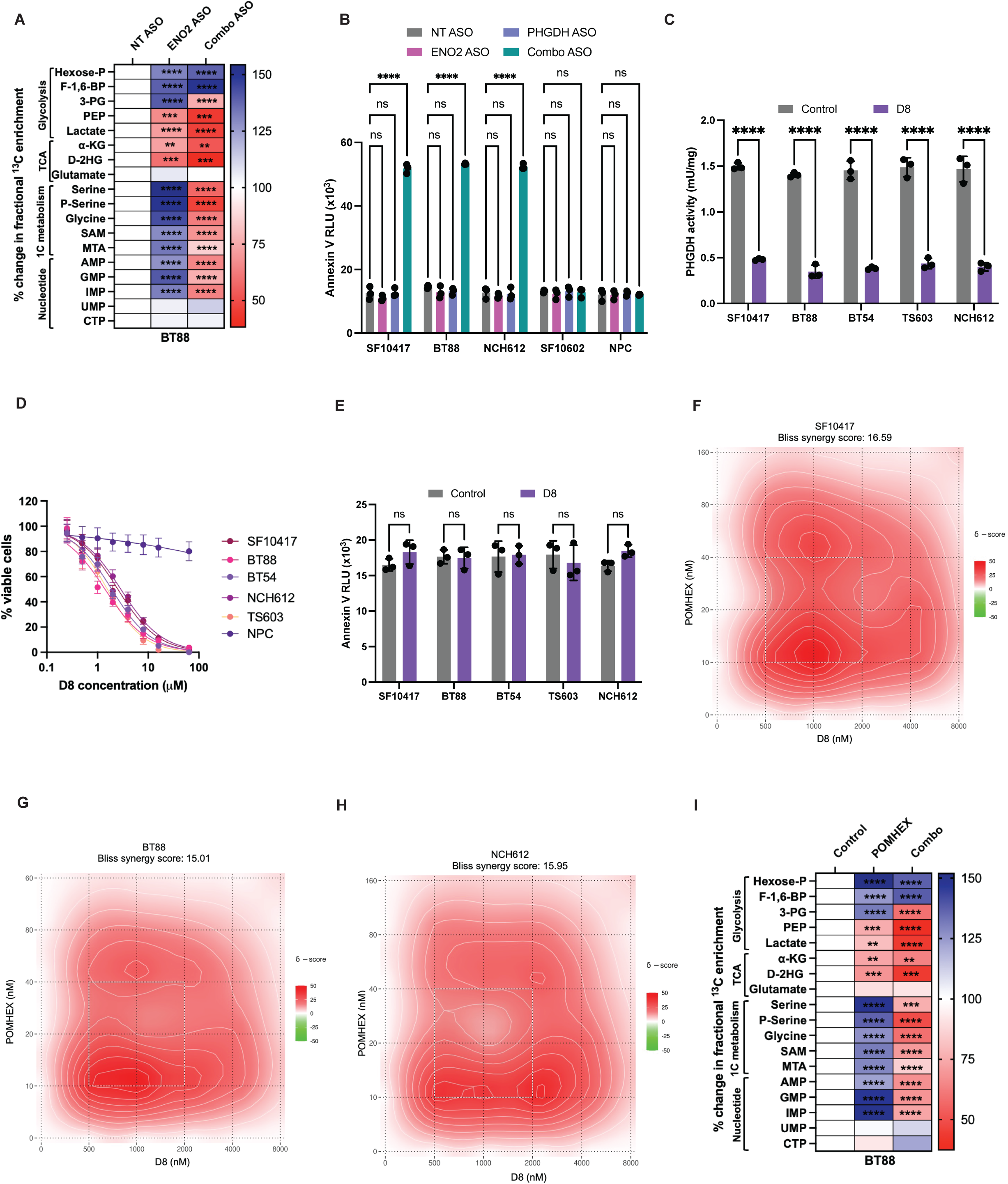
Targeting ENO2 and PHGDH is synthetically lethal in patient-derived ODG cells. **(A)** Effect of silencing ENO2 alone or ENO2 in combination with PHGDH (Combo) on fractional ^13^C metabolite enrichment from [U-^13^C]-glucose in BT88 cells. **(B)** Effect of silencing ENO2 alone or ENO2 in combination with PHGDH (Combo) on apoptosis in patient-derived ODG, AC, or NPCs. **(C)** Effect of the PHGDH inhibitor D8 on PHGDH activity in patient-derived ODG cells. **(D)** Dose-response curves showing the effect of D8 on the proliferation of patient-derived ODG cells or NPCs. **(E)** Effect of D8 on apoptosis in patient-derived ODG cells. Bliss synergy score maps for the combination of POMHEX and D8 in SF10417 **(F)**, BT88 **(G)**, or NCH612 **(H)** cells. **(I)** Effect of POMHEX or the combination of POMHEX and D8 (Combo) on fractional ^13^C metabolite enrichment from [U-^13^C]-glucose in BT88 cells. Hexose-P refers to glucose-6-phosphate or fructose-6-phosphate; F-1,6-BP refers to fructose-1,6-bisphosphate; 3-PG is 3-phosphoglycerate; PEP is phosphoenolpyruvate; α-KG is α-ketoglutarate; P-serine is phosphoserine; SAM is S-adenosylmethionine; MTA is methylthioadenosine. For panels A and I, statistical significance is represented for ENO2 inhibition vs. control and combined ENO2 and PHGDH inhibition vs. control.

4-(N-(3-Acetylphenyl) sulfamoyl)-N-(4-(3-fluorophenyl) thiazol-2-yl) benzamide (hereafter referred to as D8) was recently identified as a novel PHGDH inhibitor^41^. We confirmed that D8 inhibited PHGDH activity and abrogated the proliferation of ODG cells with IC50 values of ∼1.4-2.8 μM (Fig. 5C-5D). D8 did not inhibit the viability of NPCs (Fig. 5D). Consistent with the results of PHGDH silencing, D8 as monotherapy did not induce apoptosis of ODG cells (Fig. 5E). Importantly, the combination of POMHEX and D8 was synergistically lethal with Bliss synergy scores of 16.59, 15.01, and 15.95 for the SF10417, BT88, and NCH612 models, respectively (>10 indicates synergy; Fig. 5F-5H). Interrogation of [U-^13^C]-glucose metabolism showed that the combination of POMHEX and D8 significantly downregulated ^13^C labeling of lactate, α-ketoglutarate, D-2HG, serine, phosphoserine, glycine, S-adenosylmethionine, methylthioadenosine, AMP, and GMP in BT88 cells (Fig. 5I), consistent with the loss of both ENO2 and PHGDH activity. Taken together, these results indicate that combined inhibition of ENO2 and PHGDH abrogates glycolysis, the TCA cycle, serine biosynthesis, one-carbon metabolism, and purine nucleotide biosynthesis, leading to synergistic cell death in patient-derived ODG cells.

### The combination of POMHEX and D8 abrogates [U-^13^C]-glucose metabolism and induces tumor regression in mice bearing intracranial ODG xenografts

To validate our results *in vivo,* we examined the effect of treatment with POMHEX or the combination of POMHEX and D8 on [U-^13^C]-glucose metabolism in mice bearing intracranial patient-derived SF10417 or BT88 xenografts. Tumor-bearing mice were treated with vehicle, POMHEX, or the combination of POMHEX and D8 for 5±2 days, infused with [U-^13^C]-glucose, euthanized, and tumor tissue isolated for quantification of ^13^C metabolite labeling and correlative assays (Fig. 6A). As shown in Fig. 6B-6C, POMHEX downregulated *in vivo* glucose metabolism to lactate, α-ketoglutarate, and D-2HG, while upregulating serine, one-carbon, and purine nucleotide biosynthesis in both SF10417 and BT88 tumors. Importantly, the combination of POMHEX and D8 abrogated ^13^C enrichment of lactate, α-ketoglutarate, D-2HG, serine, phosphoserine, glycine, S-adenosylmethionine, methylthioadenosine, AMP, and GMP in mice bearing intracranial SF10417 or BT88 xenografts (Fig. 6B-6C). These metabolic alterations were associated with on-target reduction in enolase activity and upregulation of PHGDH in POMHEX-treated tumors (Fig. 6D-6E). In contrast, both enolase and PHGDH were downregulated in tumors treated with the combination of POMHEX and D8 (Fig. 6D-6E). Notably, flow cytometry for annexin V+ cells confirmed that the combination of POMHEX and D8 massively induced apoptosis, unlike monotherapy with POMHEX (Fig. 6F-6G), thereby confirming the *in vivo* synthetic lethality of the combination.

**Figure 6.**
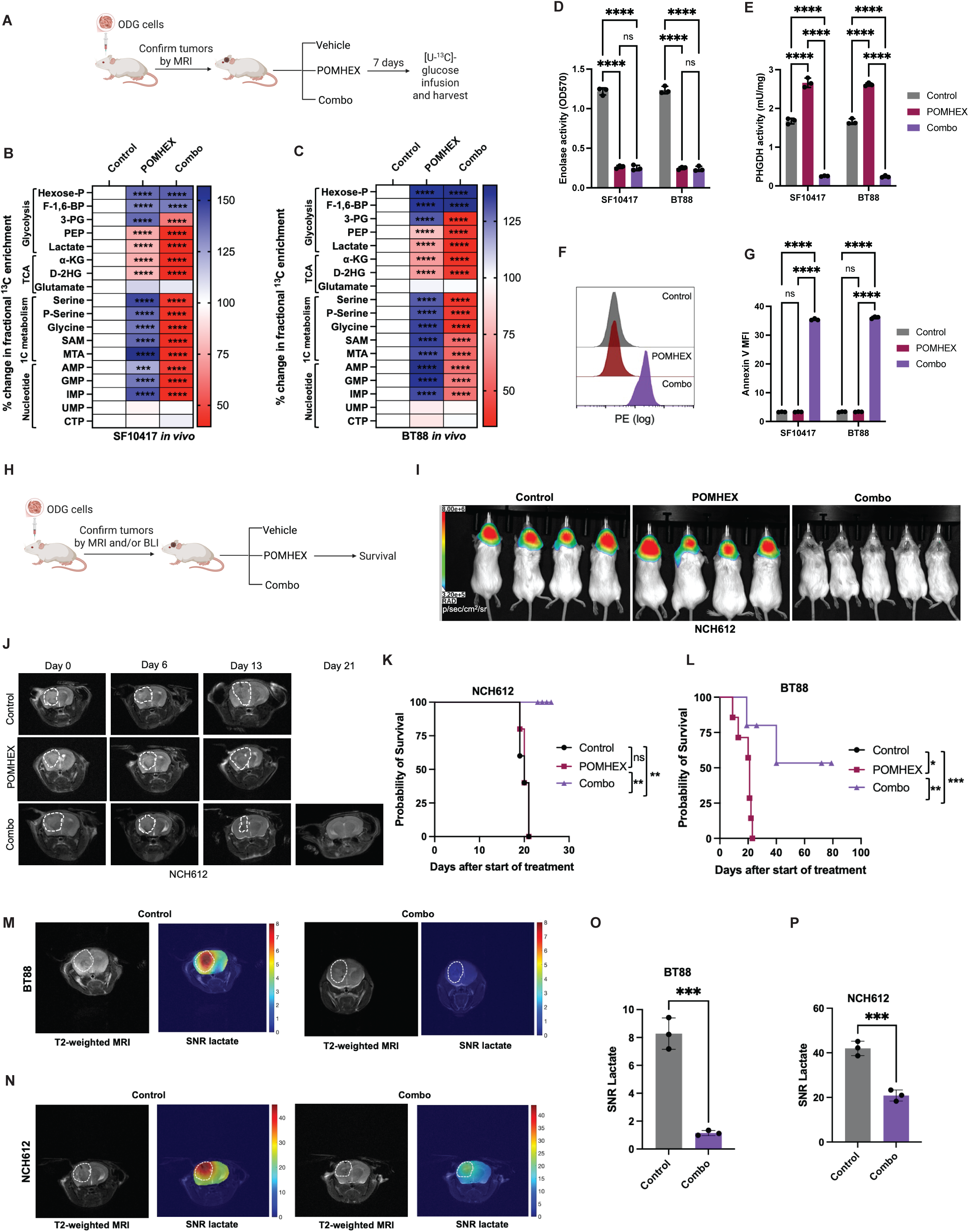
Combined inhibition of ENO2 and PHGDH induces tumor regression in mice bearing intracranial patient-derived ODG xenografts. **(A)** Schematic illustration of the study design with mice bearing intracranial SF10417 or BT88 tumors. Tumor-bearing mice were treated with vehicle, POMHEX, or the combination of POMHEX and D8 (Combo) for 7 days, infused with [U-^13^C]-glucose, and tumor tissue resected for LC-MS and correlative assays. Effect of POMHEX or the combination of POMHEX and D8 (Combo) on fractional ^13^C metabolite enrichment from [U-^13^C]-glucose in mice bearing intracranial SF10417 **(B)** or BT88 **(C)** tumors. Enolase activity **(D)** and PHGDH activity **(E)** in tumor tissue resected from mice bearing intracranial SF10417 or BT88 tumors treated with vehicle (Control), POMHEX, or the combination of POMHEX and D8 (Combo) for 7 days. **(F)** Representative flow cytometric histograms for annexin V in SF10417 tumors treated with vehicle (Control), POMHEX, or the combination of POMHEX and D8 (Combo) for 7 days. **(G)** Quantification of annexin V staining in tumor tissue resected from mice bearing intracranial SF10417 or BT88 tumors treated with vehicle (Control), POMHEX, or the combination of POMHEX and D8 (Combo) for 7 days. **(H)** Schematic illustration of the study design with mice bearing intracranial NCH612 or BT88 tumors. Tumor-bearing mice were treated with vehicle, POMHEX, or the combination of POMHEX and D8 (Combo) until the mice needed to be euthanized or the tumor was no longer visible on MRI or BLI. Representative BLI **(I)** and MR **(J)** images from mice bearing intracranial NCH612 tumors treated with vehicle, POMHEX, or the combination of POMHEX and D8 (Combo) as described in panel H. BLI images were acquired on day 17 after the start of treatment while MR images were acquired at days 0, 6, 13, and 21. Survival of mice bearing intracranial NCH612 **(K)** or BT88 **(L)** tumors treated with vehicle, POMHEX, or the combination of POMHEX and D8 (Combo) as described in panel H. Representative 2D CSI data of [6,6’-^2^H]-glucose metabolism acquired from mice bearing intracranial NCH612 **(M)** or BT88 **(N)** tumors at day 7 after treatment with vehicle or the combination of POMHEX and D8. Quantification of the SNR of lactate produced from [6,6’-^2^H]-glucose in BT88 **(O)** or NCH612 **(P)** mice. Data was acquired on a Bruker 3T (BT88) or 9.4T scanner (NCH612). Hexose-P refers to glucose-6-phosphate or fructose-6-phosphate; F-1,6-BP refers to fructose-1,6-bisphosphate; 3-PG is 3-phosphoglycerate; PEP is phosphoenolpyruvate; α-KG is α-ketoglutarate; P-serine is phosphoserine; SAM is S-adenosylmethionine; MTA is methylthioadenosine.

We then examined the effect of treatment on tumor growth and animal survival in mice bearing intracranial NCH612 or BT88 tumors *in vivo* (Fig. 6H). As shown in the longitudinal MRI and bioluminescence images in Fig. 6I-6L and Supplementary Fig. S5A, the combination of POMHEX and D8 induced tumor regression and significantly extended survival in mice bearing intracranial NCH612 or BT88 xenografts. While POMHEX as monotherapy did not impact survival in the NCH612 model, it had a small, albeit significant, effect on the survival of BT88 tumor-bearing mice (Fig. 6K-6L). Taken together, these studies highlight the therapeutic potential of targeting ENO2 and PHGDH in ODGs.

### DMI provides an early readout of response to combination therapy in mice bearing intracranial ODG xenografts

Finally, to assess whether DMI has the potential to provide an early readout of response to therapy, we quantified [6,6’-^2^H]-glucose metabolism in mice bearing intracranial BT88 or NCH612 tumors before and after treatment with vehicle or the combination of POMHEX and D8. As shown in Fig. 6M-6P, lactate production from [6,6’-^2^H]-glucose was significantly reduced within 5±2 days of combination therapy relative to vehicle-treated controls in both NCH612 and BT88 models. These results provide proof-of-principle evidence for the ability of DMI to non-invasively monitor early response to therapy in preclinical ODG models *in vivo*.

## DISCUSSION

The 1p/19 codeletion is a molecular hallmark of ODGs^2,6^. Since its discovery, this quintessential molecular alteration has been exploited for ODG diagnosis but not therapy^6^. We hypothesized that the loss of critical metabolic enzymes on chromosomes 1p or 19q could provide a novel therapeutic opportunity. Indeed, ODGs lack one copy of the glycolytic enzyme ENO1, located on chromosome 1p, due to the 1p/19 codeletion. Interestingly, our studies revealed that residual ENO1 expression is suppressed in ODGs due to promoter histone H3K27 trimethylation. Importantly, ENO2 expression is significantly upregulated, and targeting ENO2 by genetic or pharmacological means abrogates ODG viability. Furthermore, this therapeutic effect was specific to ODGs vs. ACs or NPCs, thereby identifying an opportunity for precision therapy. Of note, this concept of “collateral lethality” has previously been applied to approximately 1-5% of glioblastomas that lose ENO1 due to spontaneous loss of the 1p36 locus^11^. Our studies extend this concept to all ODGs, which, by definition, undergo 1p loss. They also suggest that the isoforms of essential metabolic genes located on chromosomes 1p or 19q, such as H6PD or CTPS2, may be druggable therapeutic targets in ODGs.

*CIC*, which is located on chromosome 19q, is the most frequently mutated gene in ODGs^6,42^. Loss of *CIC* function is associated with poorer survival in ODG patients, suggesting that CIC functions as a tumor suppressor^42^. To the best of our knowledge, we are the first to identify a role for *CIC* in regulating ENO2 expression in tumor cells. CIC functions as a transcriptional repressor downstream of the MAPK pathway in flies and mammalian cells^35^. Our data and prior studies indicate that the MAPK pathway is active in ODGs^5,7,34^. Activated MAPK signaling drives ERK1-mediated phosphorylation of CIC at S173, leading to cytosolic sequestration and loss of nuclear localization of CIC in ODG cells. The net result is derepression of ENO2 expression in ODGs. Since metabolic reprogramming fuels tumor proliferation^43^, the suppression of glycolysis via repression of ENO2 expression could be a key component of the tumor suppressive function of CIC.

Our studies are the first to identify PHGDH as a metabolic vulnerability in the context of ENO2 inhibition in cancer. Using stable isotope tracing and LC-MS to quantify [U-^13^C]-glucose metabolism, we show that genetic or pharmacological ablation of ENO2 blocks glycolysis and the TCA cycle in ODG cells and *in vivo* in mice bearing intracranial ODG xenografts. However, it redirects glucose towards the synthesis of serine, one-carbon metabolites, and purine nucleotides, an effect that is driven by upregulation of PHGDH. The resulting nucleotide imbalance blocks cell cycle progression in the S phase and inhibits viability, without significant cytotoxicity^44^. Importantly, targeting ENO2 and PHGDH in combination is synthetically lethal and leads to tumor shrinkage and extended survival of mice bearing intracranial ODG tumors *in vivo*.

Both POMHEX and D8 have the potential for clinical translation. POMHEX is a pivaloyloxymethyl (POM) pro-drug of the ENO2-selective inhibitor HEX that was designed to be cell- and blood-brain barrier (BBB)-permeable^10,11^. Our results showing that POMHEX inhibits enolase activity and abrogates [U-^13^C]-glucose metabolism downstream of enolase in mice bearing intracranial SF10417 and BT88 tumors provide pharmacodynamic evidence of BBB penetrance and on-target inhibition of glycolysis *in vivo.* Notably, POMHEX has an acceptable safety profile in rodents and non-human primates, and ENO2 knockout mice are viable^10,11^. D8 was recently developed as an orally bioavailable, competitive PHGDH inhibitor with anti-tumor activity in colorectal cancer models^41^. Our results, for the first time, indicate that PHGDH activity and [U-^13^C]-glucose metabolism to serine are abrogated in intracranial ODG tumors treated with the combination of POMHEX and D8, establishing the BBB penetrance of D8. Importantly, our studies set the stage for the exploration of D8 as a therapeutic agent for brain metastases that are also highly dependent on PHGDH for serine synthesis^45^.

Once a novel therapy has been developed, it becomes important to identify biomarkers of response to therapy^46,47^. Due to the diffusely infiltrative nature and anatomical location of brain tumors, MR-based imaging methods that can be incorporated into standard imaging protocols are desirable^47,48^. Our studies in patient-derived ODG models indicate that tracing lactate production from [6,6’-^2^H]-glucose by DMI provides an early readout of response to therapy that precedes MRI-detectable volumetric alterations and predicts extended survival *in vivo.* DMI is a clinical stage method^19,22,24^, and recent studies suggest that DMI can be interleaved with standard imaging protocols^49^. Our findings, therefore, set the stage for the incorporation of DMI into future clinical trials to assess early response to therapy in ODGs.

In summary, we have identified ENO2 as a metabolic vulnerability induced by the 1p/19q codeletion in ODGs. While targeting ENO2 as monotherapy inhibits glycolysis, it shunts glucose towards serine and purine nucleotide biosynthesis, an effect that is driven by PHGDH. Importantly, targeting both ENO2 and PHGDH is synthetically lethal and identifies a unique therapeutic opportunity for ODGs. Furthermore, visualizing glucose metabolism by DMI provides an early readout of response to therapy *in vivo*. Collectively, we have identified an integrated metabolic therapy and imaging strategy for ODGs.

## Acknowledgments

This study was funded by the National Institutes of Health grant R21CA277325 (PV) and the Gianna Rae Meadows Fund for the Oligodendroglioma Cure (PV). The authors acknowledge support from the Preclinical MR Imaging Core of the Department of Radiology and Biomedical Imaging at UCSF.

