## Supplementary methods, figures and legends for "The 1p/19q co-deletion induces targetable and imageable vulnerabilities in glucose metabolism in oligodendrogliomas"

***Cell culture:*** BT54 (female, grade 3 ODG), BT88 (male, grade 3 ODG), and BT142 (male, grade 3, AC) cells were a kind gift from Dr. Luchman from the University of Calgary, Alberta, Canada. The SF10417 (male, grade 3 ODG) and SF10602 (female, grade 3 AC) were a kind gift from Dr. Joseph Costello from the University of California, San Francisco. The NCH612 cell line (male, grade 3 ODG) was a kind gift from Dr. Christel Herold-Mende from Heidelberg University. TS603 cells were a kind gift from Dr. Sriram Venneti, University of Michigan, Ann Arbor. Human neural progenitor cells (NPCs) were purchased from ATCC (ACS-5006).

BT54 and BT142 cells were grown as neurospheres in Neurocult medium (Stem Cell Technologies) supplemented with 20 ng/mL each of human EGF and FGF2, 2 mg/mL heparin sulfate, and 1% penicillin/streptomycin^1-4^. SF10417 cells were grown as monolayers in laminin-coated flasks, and NCH612 and TS603 cells as neurospheres in serum-free Neurobasal A medium containing 2 mM glutamine, 1% penicillin/streptomycin, 2% B-27, 1% N2 supplement, 20 ng/mL each of EGF, FGF2, and PDGF-AA^5-7^. NPCs and BT88 cells were maintained as monolayers in Dulbecco’s Modified Eagle’s medium supplemented with 2 mM glutamine, 10% fetal calf serum, and 1% penicillin/streptomycin. Cell lines were routinely tested for mycoplasma contamination, authenticated by short tandem repeat fingerprinting, and assayed within 6 months of authentication. All drugs used in this manuscript were purchased from Medchemexpress.

***Quantitative PCR (QPCR):*** Gene expression was measured by QPCR using Taqman primers according to standard protocols. Briefly, RNA was isolated using the RNeasy Kit (Qiagen), and the SYBR Green QPCR kit (Sigma) was used for quantitative real-time PCR on 10 ng of cDNA template in a final volume of 20 μL. Data analysis was performed using the DDCt method with β-actin as a reference/control gene.

***Gene silencing and overexpression:*** ENO2 and PHGDH were silenced in ODG cells using AUM*silence^TM^* antisense oligonucleotides (ASOs) harboring a phosphorothioate backbone and next-generation sugar nucleotide modifications that enhance stability and enable efficient cellular uptake without the need for transfection reagents^8^. ASO sequences were as follows: ENO2 ASO#1: AGTGACCGAGCCGATCTGGTT; ENO2 ASO #2: GTGTATGGTAGACCTCTGCAC; PHGDH ASO: CCTCTTTGCTAAGGTTCTGC; NT: CCTTCCCTGAAGGTTCCTCC. The ASOs were added to the cell culture medium at a concentration of 5 μM. Silencing was confirmed by QPCR at 72 h. Two non-overlapping Accell siRNA sequences (Dharmacon) were used to silence MEK1 (encoded by *MAP2K1;* A-003571-14-0050, A-003571-15-0050), ERK1 (encoded by *MAPK1;* A-003555-19-0050, A-003555-20-0050), or AMPK (encoded by PRKAA; A-005027-14-0050, A-005027-17-0050) in ODG cells.

**Assays:** Enolase and PHGDH activity were measured using Abcam kits (ab241024 for enolase and ab273328 for PHGDH). AMPK activation was measured by using T172 phosphorylation as a marker of activation by western blotting as described above. To assay the effect of signaling pathway inhibitors, 10,000 cells/well were seeded in a 96-well plate and treated with the indicated drugs at a single concentration of 10 μM for 72 h. The Cell Lytic Buffer (Sigma) was added to the 96-well plate and incubated on ice for 30 min. Following centrifugation, the supernatant was transferred for quantification of ENO2 expression by ELISA (Proteintech, KE00050). Cytosolic and nuclear fractions were separated using the NE-PER fractionation kit (Invitrogen, 78835).

***ChIP-QPCR:*** Chromatin was immunoprecipitated using an antibody specific to H3K27me3 (Active Motif, #39155) or rabbit IgG (Cell Signaling, #2729) and the High-Sensitivity ChIP Kit (Abcam, ab185913) according to the manufacturer’s instructions. QPCR was then performed for ENO1, and the data were expressed as fold enrichment relative to IgG control.

***Immunoprecipitation:*** ~1 x 10^7^ cells or ~10 mg of tissue was lysed in RIPA buffer (25mM Tris-HCl pH 7.6, 150 mM NaCl, 1% NP-40, 1% sodium deoxycholate, 0.1% SDS) containing 0.5 mM phenylmethyl sulphonyl fluoride, 150 nM aprotinin and 1 μM each of leupeptin and E64 protease inhibitor. Lysates were cleared by centrifugation at 14,000 rpm for 15 min at 4 °C. The supernatant was pre-cleared with protein G agarose beads (Santa Cruz Biotechnology), incubated with anti-CIC antibody (Invitrogen, PA5-83721) or isotype IgG1 control (Invitrogen, MA5-47880) overnight at 4°C, followed by incubation with protein G agarose beads at 4°C for 2 h. Beads were washed 5 times with phosphate-buffered saline (PBS), and bound proteins were eluted by boiling in SDS-PAGE sample buffer (95°C for 10 min). Immunoprecipitated proteins (IP) and original lysate (input) were examined by immunoblotting as described below.

***Western blotting:*** Cells (~10^7^) or tissue (~10 mg) were lysed by incubation in Cell Lytic buffer (Sigma) containing protease inhibitors at 4°C for 20 min. Lysates were cleared by centrifugation and boiled at 95°C for 10 min in Laemmli buffer. Total protein (~20 µg) was separated on a 4-20% polyacrylamide gel by SDS-PAGE and transferred onto Immobilon-FL PVDF membranes for western blotting. Following overnight blocking in blocking buffer (ThermoFisher Scientific) at 4°C, membranes were washed and incubated with primary antibodies diluted in blocking buffer for 1 h at room temperature. Membranes were washed and treated with HRP-conjugated secondary antibodies diluted in a blocking buffer for 1 h at room temperature. After washing, the membranes were developed using an enhanced chemiluminescence substrate using an automated imager (Chemisolo; Azure Biosciences). The primary antibodies used in the study are: Phospho-AMPKα-T172 (Cell Signaling, 2575), total AMPKα (Cell Signaling, 5831), ENO1 (Abcam, ab155102), ENO2 (Agilent, IR61261-2), PHGDH (Sigma-Aldrich, HPA021241), FLAG (Sigma-Aldrich, F3165), CIC (Invitrogen, PA5-83721), Pan-phosphoserine (Invitrogen, MA1-91608), phospho-ERK1 (Cell Signaling, 4370), phospho-AKT (Cell Signaling, 4060), total ERK1 (Cell Signaling, 9102), total AKT (Cell Signaling, 4691), GAPDH (Cell Signaling, 2118), cyclophilin B (Cell Signaling, 43603), β-actin (Cell Signaling, 4970), and goat anti-rabbit IgG-HRP (Cell Signaling, 7074).

***Flow cytometry:*** Cell cycle progression was measured by staining live cells for DNA content using the Vybrant™ DyeCycle™ Green stain (Thermo Scientific). 1x10^6^ cells were stained with 10 μM Vybrant™ DyeCycle™ Green and incubated in the dark at 37°C for 30 min. Cells were analyzed by flow cytometry on a MACSQuant 10 analyzer with excitation at 488 nm and emission at 525 nm.

To measure doubling time, cells were seeded in a 96-well plate at 10,000 cells per well and treated as described above. After 4 days, cell number was measured using the MACSQuant 10 analyzer. The doubling time was calculated using the following formula: doubling time = [4 days × (ln2)] / [ln (day 4 cell count/day 0 cell count)].

To quantify protein expression, cells were fixed and permeabilized by resuspension in the Cytofix/Cytoperm fixation solution (BD Biosciences) followed by incubation in the dark at 4°C for 20 min. Cells were washed and intracellularly stained with the relevant antibody at the dilution recommended by the manufacturer for 20 min at room temperature. Mean fluorescence intensity was measured using a MACSQuant 10 flow cytometer. The antibodies used were: ENO1 (Proteintech, CL488-11204), ENO2 (Proteintech, CL488-66150), phospho-ERK1 (Invitrogen, 46-9109-42), phospho-AKT (Invitrogen, 25-9715-42), CIC (Invitrogen, PA5-83721) and pan-phosphoserine (Invitrogen, MA1-91608).

To quantify apoptosis in BT88 and SF10417 tumors, tumor tissue was dissociated into single cells using the tumor dissociation kit (Miltenyi Biotec) according to the manufacturer's instructions. Mouse cells were then depleted using the mouse cell depletion kit (Miltenyi Biotec) according to the manufacturer's instructions. cells were resuspended in annexin binding buffer and stained with a 1:50 dilution of the anti-annexin V-PE antibody (BD Biosciences, 556421) for 30 min in the dark at room temperature. Cells were washed once in the annexin binding buffer, and the % of annexin V+ cells was measured using a MACSQuant 10 flow cytometer.

***PLA:*** The Duolink flowPLA Mouse/Rabbit Starter Kit Far Red (Sigma-Aldrich, DUO94104) was used according to the manufacturer’s instructions^9^. Briefly, cells were treated as needed and fixed and permeabilized by resuspension in the Cytofix/Cytoperm solution in the dark at 4°C for 20 min. Cells were washed once with Duolink wash buffer and resuspended in Duolink blocking solution at a concentration of 1x10^6^ cells in 100 μL. Following incubation at 37°C for 1 h, cells were pelleted and resuspended in 100 μL of Duolink antibody diluent containing the rabbit anti-CIC antibody (1:100) and the mouse anti-pan-phosphoserine antibody (1:100). For the negative isotype IgG controls, cells were resuspended in 100 μL of Duolink antibody diluent containing the mouse IgG1 isotype control (1:100; ThermoFisher Scientific, MA1-10407) and the rabbit IgG isotype control (1:100, PTMBio, PTM-5073). We also examined cells resuspended in each primary antibody alone as added controls. Following incubation at 37°C for 1 h, the cells were washed twice with Duolink wash buffer and resuspended in 100 μL of the Duolink antibody diluent containing the anti-mouse MINUS PLA probe and the anti-rabbit PLUS PLA probe at a 1:5 dilution. After incubation at 37°C for 1 h, the cells were washed twice with Duolink wash buffer and resuspended in 50 μL of ligation buffer (containing two bridging DNA oligonucleotides and a ligase at a 1:40 dilution). The DNA oligonucleotides hybridize with the two PLA probes, allowing the ligase to form a rolling circle of DNA when the probes are in close proximity (for example, when the primary antibodies are bound to a protein and a post-translational modification on the protein). After incubation at 37°C for 30 minutes, the cells were washed twice in Duolink wash buffer and incubated with 50 μL of amplification solution (containing nucleotides, fluorescently labeled, complementary oligonucleotides, and DNA polymerase at a 1:80 dilution) for 100 minutes at 37°C. The DNA polymerase amplifies the circular DNA via rolling circle amplification, and the fluorescently labeled oligonucleotides hybridize to a specific sequence in the amplified DNA to yield an amplified fluorescence signal that is detectable by flow cytometry ^10-12^. The cells were then washed twice in Duolink wash buffer, resuspended in stain buffer, and analyzed on a MACSQuant 10 flow cytometer with excitation at 644 nm and emission at 669 nm.

***Stable isotope tracing in cells:*** Cells were treated as needed and incubated in media in which glucose was replaced with [U-^13^C]-glucose (99% enrichment; Cambridge Isotope Laboratories, final concentration 25 mM) for 72 h. After washing in ice-cold ammonium acetate (150 mM, pH 7.3), metabolites were extracted by the addition of 1 ml of ice-cold methanol/water (80:20 v/v). The samples were reconstituted with 60 μL of ice-cold acetonitrile/water (50:50, v/v), transferred into glass vials, and utilized for liquid chromatography-mass spectrometry (LC-MS) as described below.

***Stable isotope tracing in vivo:*** Studies were performed on mice bearing intracranial BT88 or SF10417 tumors treated with vehicle (saline), POMHEX (25 mg/kg), or the combination of POMHEX and D8 (25 mg/kg each) daily for 5 days per week via intraperitoneal injection. At day 7 post-treatment, mice were intravenously injected with a bolus of 428 mg/kg of [U-^13^C]-glucose dissolved in sterile saline, followed by continuous infusion at 14 mg/kg/min for 2 h. Tumor and contralateral normal brain tissue were collected and snap frozen. Metabolites were extracted from ~15-25 mg of tissue using 1 ml of ice-cold methanol/water (80:20 v/v), lyophilized, reconstituted in 60 μL of ice-cold acetonitrile/water (50:50, v/v), and examined by LC-MS as described below.

***LC-MS:*** LC-MS was performed using a Vanquish Ultra High-performance LC system coupled to an Orbitrap ID-X mass spectrometer (ThermoFisher Scientific)^13^ (see Supplementary Information for a detailed description). Metabolites were separated using a Luna 3 NH2 column (150 mm x 2.1 mm, 3 μm, Phenomenex) using mobile phases A (5 mM ammonium acetate, 48.5 mM ammonium hydroxide pH 9.9) and B (100% Acetonitrile). High-resolution MS was acquired using a full scan method alternating between positive and negative polarities. MS1 data were acquired at a resolution of 60,000 with a standard automatic gain control and a maximum injection time of 100 ms using the Xcalibur software. Peak areas were quantified using TraceFinder, corrected to blank samples, and normalized to the total ion count and the cell number, protein content, or wet weight of the sample. The % ^13^C enrichment was calculated after correcting for natural abundance using Escher Trace^14^.

***In vivo studies:*** All studies were approved by the University of California, San Francisco Institutional Animal Care and Use Committee (IACUC). ODG cells (5x10^5^ cells in 3 μL) were intracranially injected into the cortex of SCID mice (female, 5-6 weeks old)^15-17^. Once tumors were visible on MRI or BLI, this timepoint was considered D0, and mice were randomized and treated with vehicle (saline), POMHEX (25 mg/kg), or the combination of POMHEX and D8 (25 mg/kg each) daily for 5 days per week via intraperitoneal injection. Mice were treated until they needed to be euthanized per IACUC guidelines, or the tumor was no longer visible on MRI. Tumor volume was determined by T2-weighted MRI using a small animal horizontal Bruker 3T or 9.4T scanner^15-17^ For the 3T scanner, we used a ^1^H quadrature volume coil and a T2 rapid acquisition with relaxation enhancement (RARE) sequence (TE/TR = 64/3700 ms, FOV = 30 x 30 mm^2^, matrix = 256 x 256, slice thickness = 1.5 mm, NA = 5). Studies at 9.4T were performed using a ^1^H /^13^C linear/linear volume coil on a preclinical 9.4T MR scanner (Biospec, Bruker). Axial T2-weighted images were acquired using a spin-echo TurboRARE sequence (TE/TR = 8.25/3200ms, FOV = 30×30 mm^2^, 256×256, slice thickness=1.2 mm, NA=3). Radiance was quantified on a Spectral Instruments Imaging (ATX) after intraperitoneal injection of 150 mg/kg D-luciferin.

***DMI studies in vivo*:** DMI studies were performed on a small animal horizontal Bruker 3T or 9.4T scanner using a 16 mm ^2^H surface coil in combination with a quadrature ^1^H coil (40 mm inner diameter for the 3T scanner and 72 mm inner diameter for the 9.4T scanner). For spatial localization of lactate production, a 2D chemical shift imaging (CSI) sequence was used (3T: TE/TR = 1.04/265.89 ms, FOV = 30 x 30 x 8 mm^3^, complex points=128 points, spectral width=2.5 kHz, NA = 30, temporal resolution=8 minutes 30 s, nominal voxel size = 112.5 μL; 9.4T: TE/TR = 1.104/591.824 ms, FOV = 30 x 30 x 5 mm^3^, 512 points, spectral width = 2.5 kHz, NA = 8, temporal resolution=8 minutes 30 s, nominal voxel size = 70.3 μL. 2D CSI data was analyzed using in-house MATLAB codes as described earlier^15,16^. For each voxel at every timepoint, peak integrals were calculated. To generate heatmaps of the SNR of ^2^H-lactate, raw data were interpolated from an 8x8 matrix to a 256x256 matrix and normalized to noise. The SNR of ^2^H-lactate was quantified from a region of interest of 10.99 mm^3^ (3T) or 6.87 mm^3^ (9.4T) placed over the tumor.

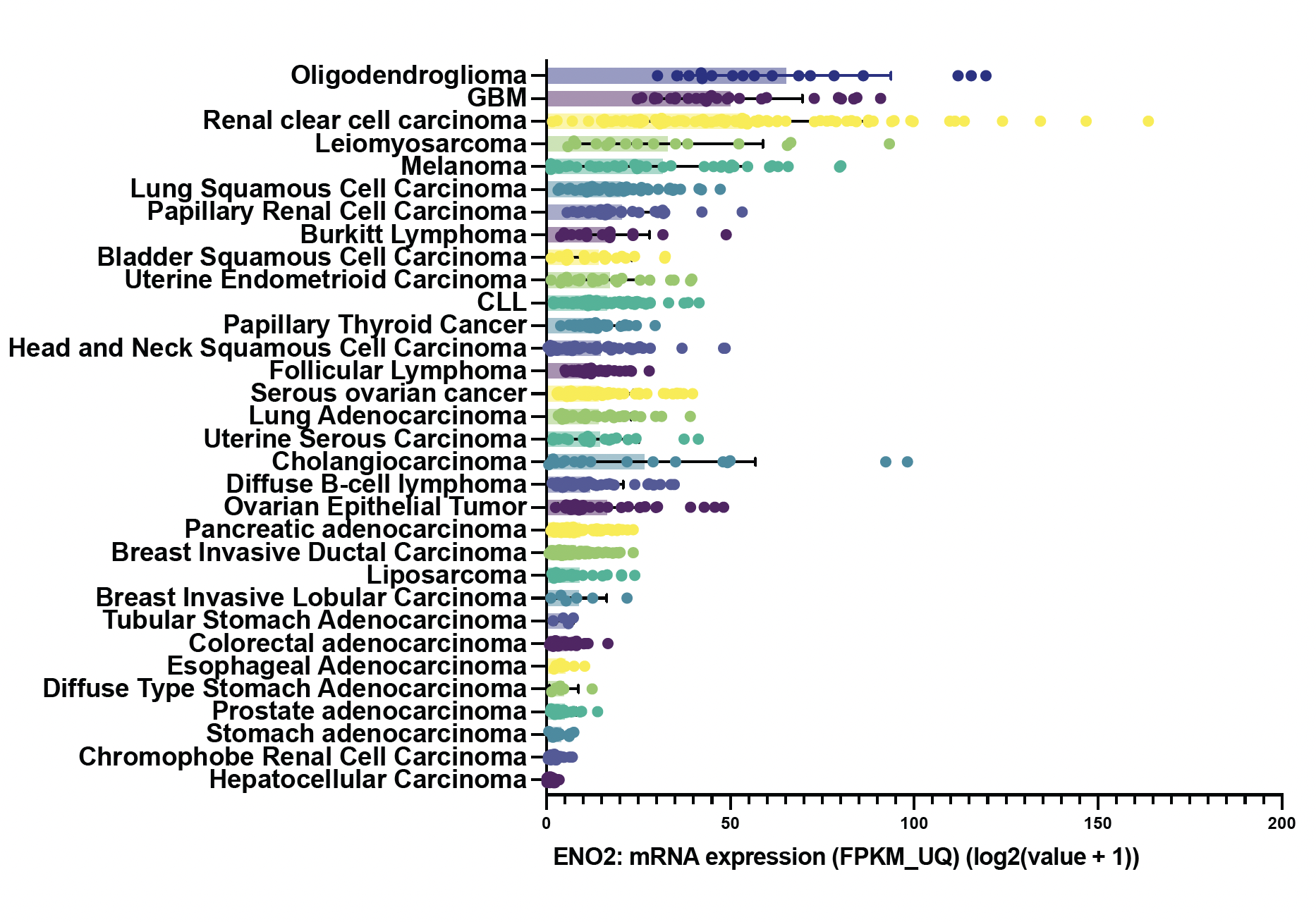
**Supplementary Figure S1. ENO2 expression in the TCGA database.** ENO2 mRNA abundance across different cancer types from the TCGA pan-cancer studies database (cBioportal).

**Supplementary Figure S2. The MAPK pathway drives ENO2 expression in ODGs.** Quantification of the mean fluorescence intensity of phospho-ERK1 **(A)** and phospho-AKT **(B)** in patient-derived ODG (SF10417, BT88, BT54, TS603, NCH612), AC (SF10602, BT142), and NPCs. Verification of MEK1 **(C)** and ERK1 **(D)** silencing in ODG cells treated with siRNAs targeting a non-specific sequence (siControl), MEK1 (siMEK1), or ERK1 (siERK1). In each case, 2 non-overlapping siRNA sequences were used. **(E)** CIC mRNA abundance in patient-derived ODG (SF10417, BT88, BT54, TS603, NCH612), AC (SF10602, BT142), and NPCs. **(F)** Quantification of CIC phosphorylation as measured by the PLA in patient-derived ODG (SF10417, BT88, BT54, TS603, NCH612), AC (SF10602, BT142), and NPCs. **(G)** Quantification of the effect of inhibiting the MAPK pathway using trametinib on CIC phosphorylation as measured by the PLA in SF10417, BT88, and NCH612 cells. **(H)** Verification of the expression of FLAG-tagged mutant S173A CIC in SF10417 cells by western blotting. β-actin was used as the loading control.

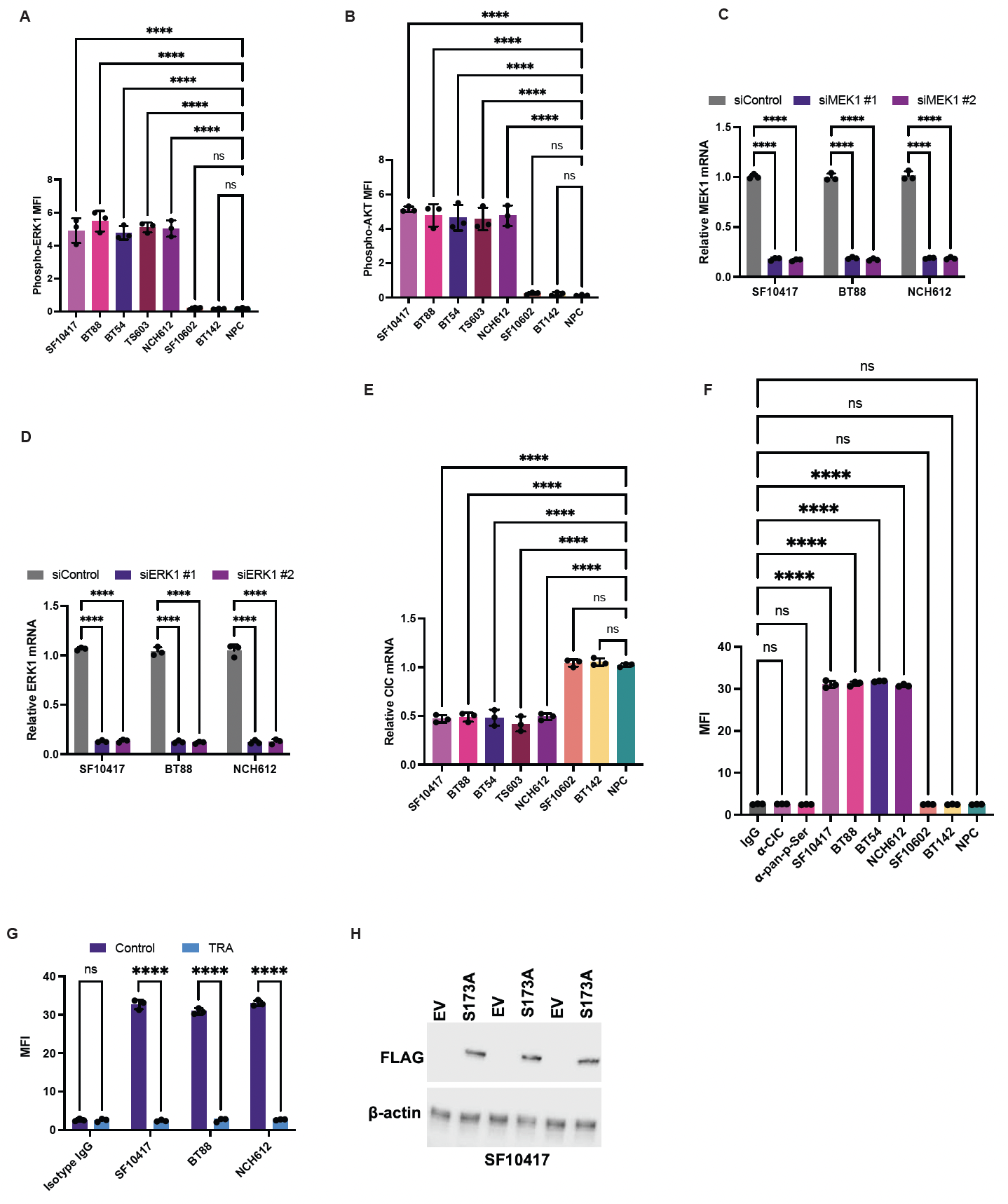
**Supplementary Figure S2.**

**Supplementary Figure S3. Effect of silencing or inhibiting ENO2 in ODGs. (A)** Verification of ENO2 silencing in ODG (SF10417, BT88, NCH612), AC (SF10602, BT142), and NPC cells treated with non-targeting ASO (NT ASO), or 2 non-overlapping ASOs against ENO2. Effect of silencing ENO2 **(B)** or POMHEX **(C)** on apoptosis as measured by annexin V in patient-derived ODG (SF10417, BT88, BT54, TS603, NCH612) cells. Effect of silencing ENO2 **(D)** or POMHEX **(E)** on ATP abundance in patient-derived ODG (SF10417, BT88, BT54, TS603, NCH612) cells. **(F)** Effect of POMHEX on the AMP/ATP ratio in patient-derived ODG (SF10417, BT88, BT54, TS603, NCH612) cells.

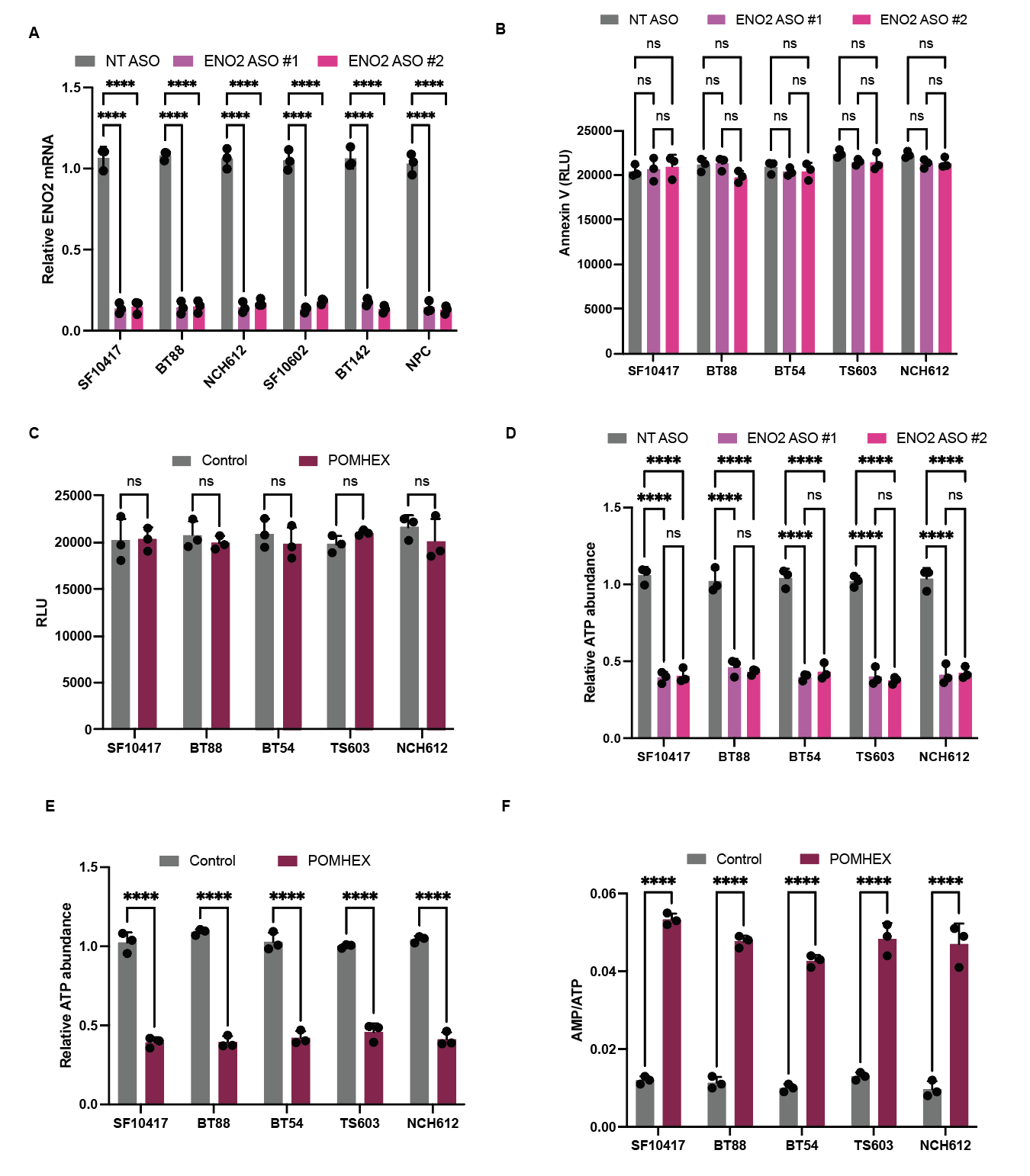
**Supplementary Figure S3.**

**Supplementary Figure S4. Effect of silencing ENO2 and PHGDH in ODGs.** Verification of ENO2 silencing **(A)** and PHGDH silencing **(B)** in SF10417, BT88, and NCH612 cells treated with NT ASOs, ENO2 ASO alone, PHGDH ASO alone, or ENO2 ASO in combination with PHGDH ASO.

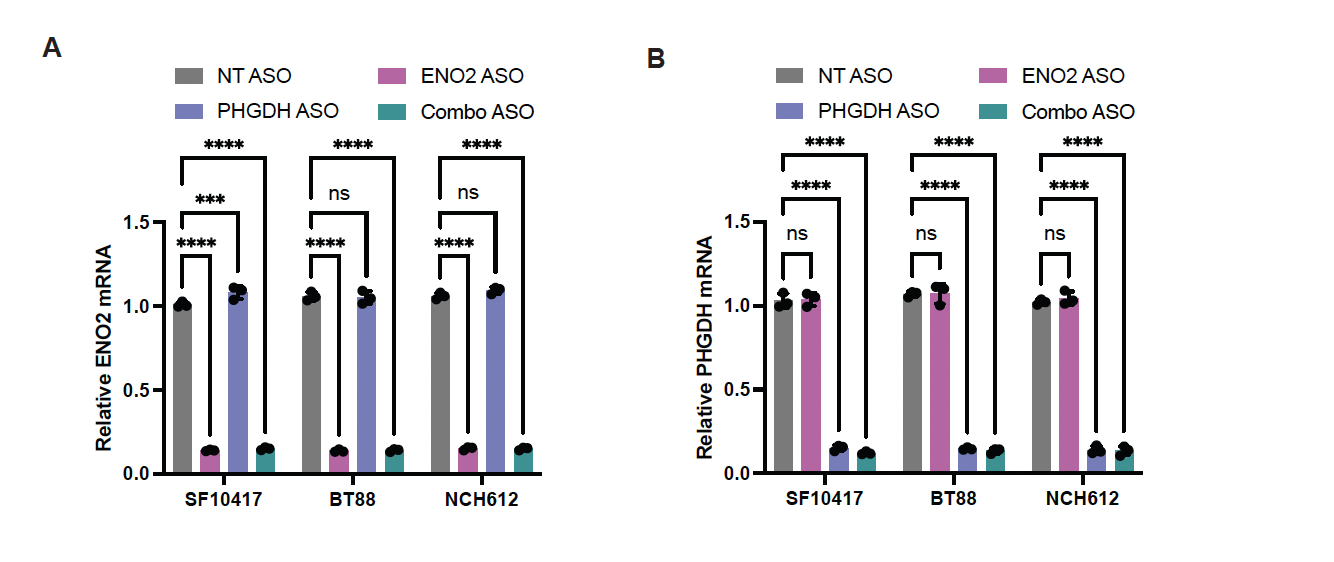

**Supplementary Figure S5. Targeting ENO2 and PHGDH in combination is synthetically lethal *in vivo* in mice bearing intracranial ODG xenografts. (A)** Effect of treatment with vehicle, POMHEX, or the combination of POMHEX and D8 on tumor growth over time as measured by BLI in mice bearing intracranial NCH612 xenografts.

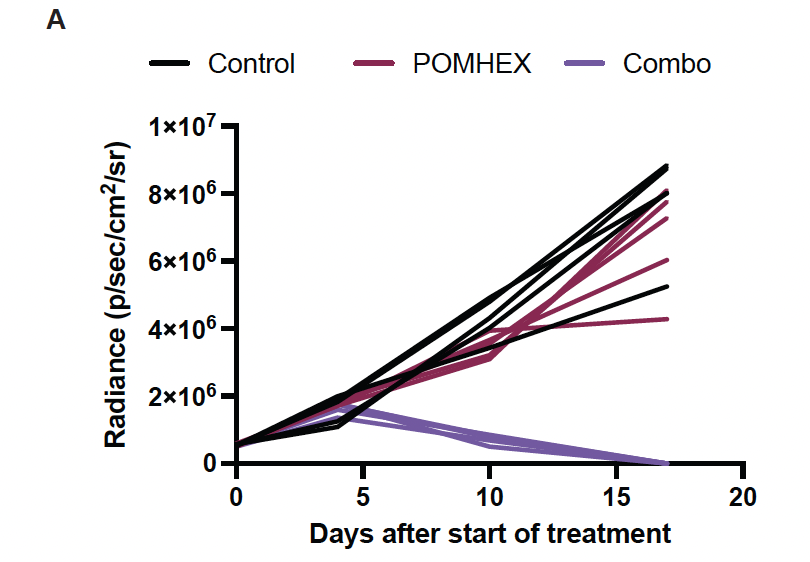
